# Small but Mighty: The Surprising Metabolic Range of a Bacterial Microcompartment

**DOI:** 10.64898/2026.06.05.730297

**Authors:** Dorian Gregnanin, Denis Jallet, Andrew W. Strong, Léa Jasse, Vanessa Soldan, Evelyne Deery, Martin Warren, Pierre Millard, Josh V. Vermaas, Stéphanie Heux

## Abstract

Bacterial microcompartments (BMCs) are protein-based metabolic modules increasingly investigated as potential intracellular nanoreactors. In this study, we engineered *Escherichia coli* by chromosomally integrating the propanediol utilization (*pdu*) operon from *Citrobacter freundii* and evaluated the metabolic versatility of the heterologously produced Pdu BMCs. The formation of well-assembled, operational Pdu BMCs was confirmed *in cellula*. Oxygen was found to inhibit 1,2-propanediol utilization, an effect alleviated under microaerobic conditions. The engineered *E*. *coli* strain assimilated a range of diols and triols, demonstrating that Pdu BMCs process both their native substrate and glycerol and linear diols up to 1,2-butanediol (C4). This substrate promiscuity was exploited to produce high value-added compounds and revealed previously unrecognized potential for plastic upcycling through ethylene glycol metabolism. In keeping with molecular dynamics simulations indicating that the Pdu BMC shell does not impose significant diffusion barriers for longer chain diols and triols, the range of amenable substrates was successfully extended to 1,2-pentanediol (C5) by mutating the diol dehydratase large subunit (PduC). Finally, deletion of *pduPQ* revealed that host metabolism partially compensated for the loss of BMC-associated aldehyde-processing enzymes, but did not reach the conversion efficiency and product distribution supported by the intact Pdu system. Together, these findings establish heterologous Pdu BMCs as promiscuous and genetically tunable metabolic modules in *E*. *coli*, highlighting their potential as scalable platforms for synthetic biology and biotechnological applications.

## Introduction

Bacterial microcompartments (BMCs) are large macromolecular assemblies consisting of a selectively permeable protein shell encapsulating particular metabolic pathways ^1^. These structures are found in the cytosol of approximately 20% of bacterial species ^2,3^ and confer a selective advantage in certain ecological niches by enhancing metabolic flexibility. BMCs are generally associated with the metabolism of particular substrates; for example, propanediol utilization (Pdu) BMCs enable the assimilation of 1,2-propanediol (1,2-PDO). Pdu BMCs are encoded by the chromosomal PDU locus, which comprises a central operon flanked by ancillary genes ^3^. This locus is widely distributed across bacterial taxa, including *Citrobacter freundii*, *Salmonella enterica*, *Listeria monocytogenes*, *Klebsiella* spp., *Clostridium* spp. as well as *Lactobacillus* spp.^2^. In Enterobacteriaceae, the *pdu* operon typically spans 19 kilobases and contains 19-21 genes encoding BMC shell proteins, pathway enzymes and enzymes required for the regeneration of cofactors ^5^. Expression of the *pdu* operon is regulated by PocR, an AraC/XylS family transcription factor that activates the P*_pdu_* promoter in response to 1,2-PDO ^6–8^.

The PDU locus generally includes a transporter (e.g. PduF) that enables uptake of extracellular 1,2-PDO. Once internalized, 1,2-PDO enters into the BMC, where it is converted into propionaldehyde by the multienzyme complex propanediol dehydratase (PduCDE) ^9^. Propionaldehyde is a reactive and toxic intermediate that can damage proteins and nucleic acids, leading to cellular stress ^9^. Encapsulation within the BMC is thought to mitigate this toxicity by sequestering the intermediate and promoting efficient downstream conversion ^10^. Within the compartment, propionaldehyde is metabolized via two competing branches: a reductive pathway yielding 1-propanol and an oxidative pathway producing propionyl-CoA and, ultimately, propionate. The reductive branch, catalyzed by an alcohol dehydrogenase (PduQ), primarily serves to regenerate NAD⁺ for redox balance ^11^, whereas the oxidative branch, mediated by propionaldehyde dehydrogenase (PduP), phosphotransacylase (PduL) and propionate kinase (PduW), generates NADH and ATP ^5,12^ (Fig. 1A). Propionate can subsequently be assimilated into central metabolism, via the methylcitrate cycle for example ^13^. As well as mitigating toxicity, BMCs enhance metabolic efficiency by colocalizing enzymes and substrates in a confined microenvironment, as well as by promoting substrate channeling, increasing local intermediate concentrations and reducing flux through competing cytosolic pathways ^14^.

**Figure 1:**
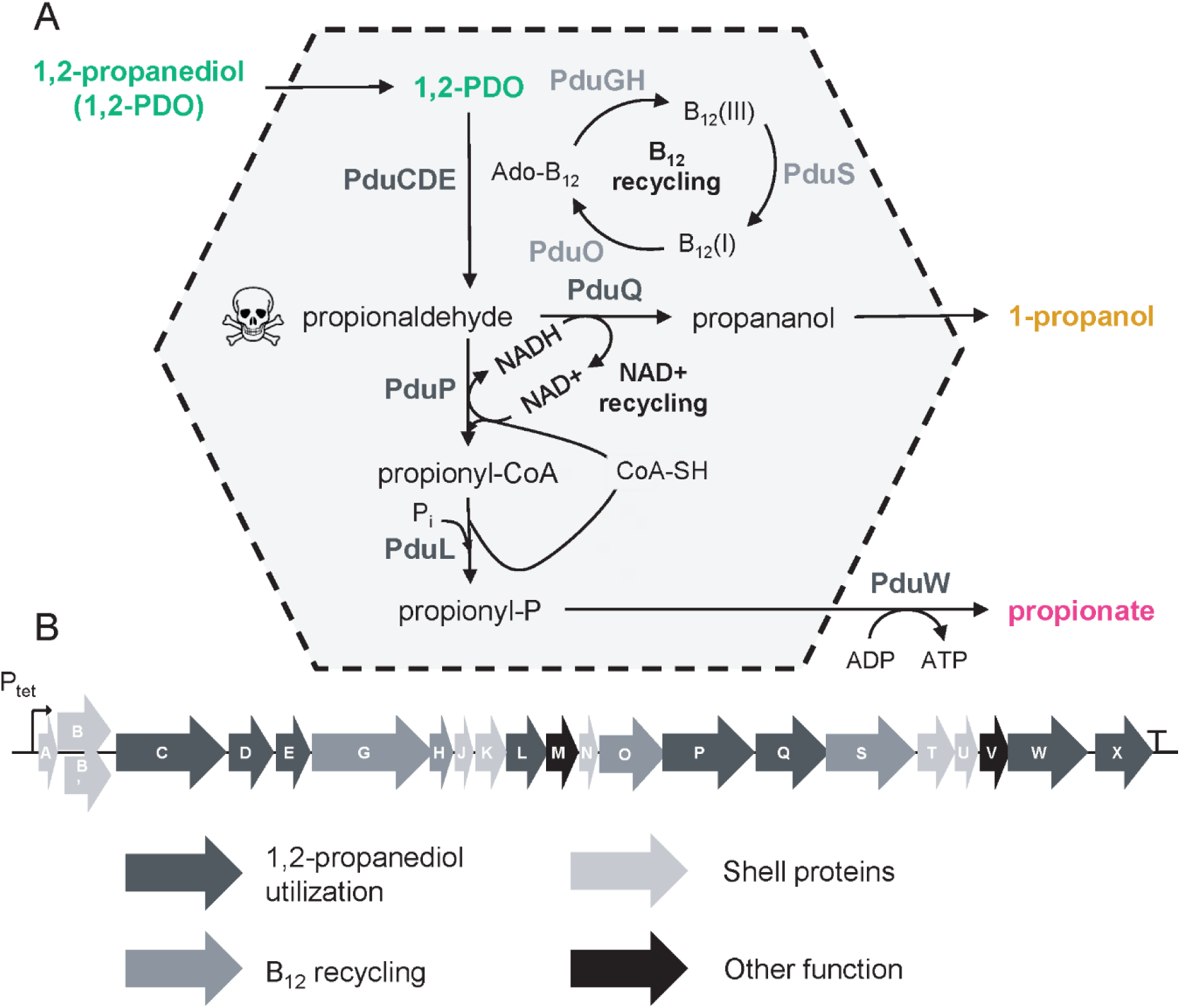
The Pdu BMC of *Citrobacter freundii*. (A) The Pdu BMC shell encapsulates enzymes required for 1,2-propanediol (1,2-PDO) conversion into 1-propanol and propionate. (B) The *pdu* operon from *Citrobacter freundii* was integrated into the chromosome of *Escherichia coli* K-12 BW25113. The *pdu* operon encodes structural proteins (PduABB’JKNTU), catalytic enzymes involved in 1,2-PDO degradation (PduCDELPQW), B₁₂ recycling proteins (PduGHOSX) and additional proteins with other proposed function. PduV has been proposed to mediate filament-associated BMC movement ^60^ while PduM is thought to act as a linker between the shell interior and the enzymatic core ^61^.

The Pdu BMC shell is widely considered to be selectively permeable, facilitating the passage of small metabolites such as 1,2-PDO, 1-propanol and propionate but restricting the diffusion of propionaldehyde and other reactive intermediates ^15,16^. This selectivity is mediated by shell protein pores, whose size, electrostatic properties and conformational dynamics govern molecular transport. Although results suggest that permeability is regulated at several levels, the underlying mechanisms remain incompletely understood ^17^. Larger cofactors required for enzymatic activity within the BMC lumen may also pass through the shell via specialized or gated pores ^18,19^. Furthermore, it has been proposed that the internal enzyme PduS may reoxidize NADH and transfer electrons through an iron-sulfur cluster in the shell protein PduT to an as-yet unidentified cytosolic acceptor ^20–22^.

Although Pdu BMCs are classically associated with 1,2-PDO metabolism, growing evidence suggests a broader substrate scope. In *Lactobacillus reuteri, Salmonella enterica* and *Klebsiella pneumoniae*, glycerol is converted into 3-hydroxypropionaldehyde (3-HPA), 1,3-propanediol (1,3-PDO), and 3-hydroxypropionate (3-HP) through Pdu-associated pathways ^23–25^. *In vitro* studies further showed that the PduCDE dehydratase can process glycerol and several short-chain 1,2-diols, while PduP and PduQ accept a wider range of aldehydes ^26^. Consistently, *pdu* genes are induced by substrates such as ethylene glycol and 2,3-butanediol in diverse bacteria ^27–29^, and the regulator PocR responds to several diols besides 1,2-PDO ^30^. Together, these observations suggest that Pdu BMCs are more metabolically versatile than previously appreciated, although this potential has not been systematically investigated.

One strategy to assess the metabolic capabilities of Pdu BMCs in a controlled framework is heterologous expression in a tractable host. To date, however, heterologous production of complete Pdu BMCs has only been reported for the *Citrobacter freundii pdu* in *E. coli* ^22^ and for the *S*. *enterica pdu* in multiple bacterial species ^31^, with limited characterization of the metabolic scope and versatility of these systems. We address this gap by providing a detailed assessment of Pdu BMC function in a heterologous *E. coli* system. By integrating chromosomal pathway engineering, microscopy, metabolite profiling and computational modelling, we systematically evaluate native and non-native substrate utilization. Our results reveal the extent of Pdu BMC metabolic promiscuity and establish a basis for their development as tunable, modular platforms for sustainable biotechnological applications.

## Results and discussion

### Heterologous Pdu BMCs are well formed and metabolically active in E. coli

Most *E*. *coli* strains, including biotechnological chassis such as K-12, BL21 and JM109, do not have a native PDU locus. Parsons et al. (2008) demonstrated heterologous Pdu BMC formation in *E*. *coli* using a cosmid carrying the *C. freundii pdu* operon ^22^. However, in native hosts, the *pdu* operon is chromosomally encoded. To mimic this genetic context, we integrated the *C*. *freundii pdu* operon into the *cobU-yoeG* locus of the *E. coli* K-12 BW25113 chromosome (Fig. 1B; Fig.S1). To decouple *pdu* expression from its native regulatory inputs, we placed the operon under the control of a tetracycline inducible promoter (P*_tet_*), thereby eliminating the requirement for 1,2-PDO induction and enabling systematic assessment of substrate scope. Chromosomal integration into the resulting BW Pdu strain was confirmed by PCR (Fig. S2, Table 1).

**Table 1:**
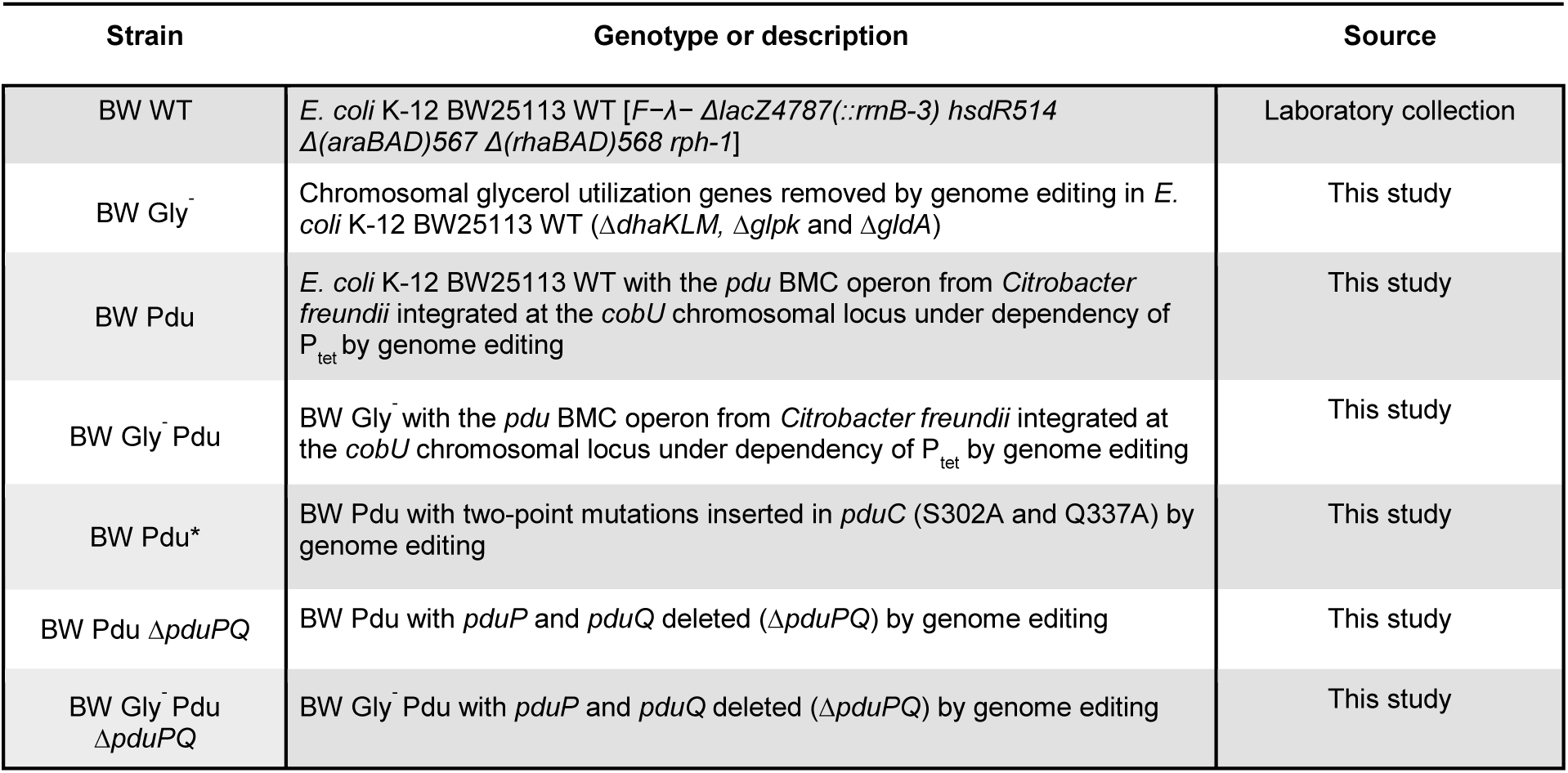
Bacterial strains used in the study.

To evaluate whether this strain produces well-formed Pdu BMCs, we purified BMCs using a multi-step centrifugation protocol after initial extraction ^32^ (see Supplementary Materials and Methods). SDS-PAGE results from the purified fraction closely matched the polypeptide profile of Pdu BMCs purified from *S*. *enterica* (Fig. 2A) ^32,33^. In contrast, the same protocol applied to the BW wild-type strain (BW WT) yielded only a diluted fraction with no visible Pdu-associated protein bands. Transmission electron microscopy (TEM) analysis of various BW WT and BW Pdu cultures showed that preservation was optimal using high-pressure fixation (HPF; see Supplementary Materials and Methods) during exponential growth in M9 minimal medium supplemented with D-glucose, 1,2-PDO, cyanocobalamin and tetracycline (Fig. 2B&C, Fig. S3). Distinct cytosolic structures roughly 100 nm in diameter were observed in BW Pdu but not in BW WT (Fig 2B&C, Fig. S3). These structures are consistent with previous observations of Pdu BMCs in cells^22^. Taken together, these results demonstrate that heterologous Pdu BMCs are correctly assembled in the BW Pdu strain.

**Figure 2:**
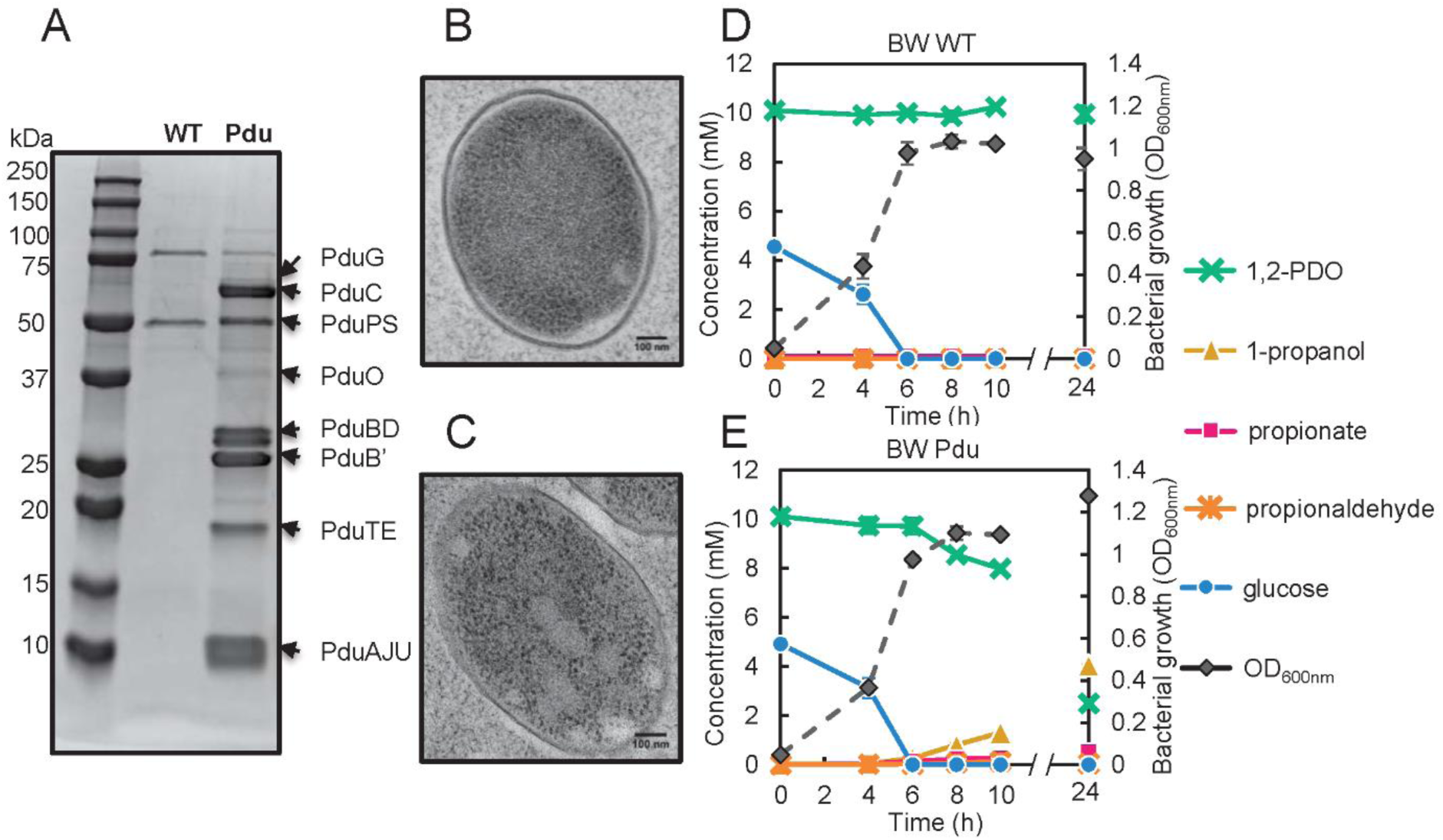
Heterologous Pdu BMCs are well formed and functional in *E*. *coli* K-12 BW25113 Pdu. (A) Comparison of SDS-PAGE separations from the purified BMC fraction of BW Pdu (Pdu) BW wild-type (WT) (See supplementary Materials and Methods). (B) and (C) Transmission electron microscopy images of (B) BW WT and (C) BW Pdu strains grown aerobically in M9 minimal medium containing D-glucose, 1,2-PDO, cyanocobalamin and tetracycline (See supplementary Materials and Methods). (D) and (E) Bacterial growth was monitored by measuring optical density at 600 nm (OD_600nm_) while substrate consumption (1,2-PDO and D-glucose) and product excretion (1-propanol and propionate) were quantified by ¹H NMR for the (D) BW WT and (E) BW Pdu strains. Data are the averages of n = 3 biological replicates and the error bars show the corresponding standard deviations

BW WT and BW Pdu strains grew at similar rates (µ = 0.40 ± 0.05 h^−1^ for BW WT and µ = 0.49 ± 0.01 h^−1^ for BW Pdu; Fig. 2D&E) in M9 medium supplemented with D-glucose, 1,2-PDO, cyanocobalamin and tetracycline under fully aerobic conditions, indicating that heterologous *pdu* expression does not impose a detectable fitness burden. Although Pdu BMCs involve thousands of protein subunits per compartment ^33^, their assembly in BW Pdu does not seem to incur a significant metabolic cost. NMR analysis of culture supernatants revealed complete deletion of D-glucose within 6 h (Fig. 2D&E), confirming its role as the primary carbon source for growth. As expected, BW WT did not consume 1,2-PDO (Fig. 2D). In contrast, BW Pdu consumed approximately 20% of the supplied 1,2-PDO in 10 h (Fig. 2E) and produced 1.30 mM 1-propanol and 0.27 mM propionate (Fig. 2E), corresponding to 61.5% and 12.7% conversion of the consumed 1,2-PDO.

Taken together, these findings demonstrate that chromosomal integration of the *pdu* operon enables the formation of functional Pdu BMCs in *E. coli*. However, under aerobic culture conditions, 1,2-PDO utilization was partial and relatively slow, indicating suboptimal pathway performance potentially influenced by environmental factors.

### Heterologous Pdu BMCs are more metabolically active under microaerobic conditions

Previous enzymatic studies have shown that the alcohol dehydrogenase PduQ is susceptible to oxygen-dependent inactivation because of its iron-containing catalytic site ^11^. We therefore investigated whether oxygen availability influences Pdu BMC function *in cellula*. BW Pdu cultures were grown in crimp-sealed vials with medium occupying 24%, 36%, 50%, 72% or 96% of the total volume, with higher filling volumes corresponding to lower oxygen availability. Cultures were shaken at 200 rpm and started without oxygen purging, allowing initial aerobic growth, followed by gradual establishment of microaerobic conditions as oxygen was consumed.

Glucose was consumed before 1,2-PDO in all cases (Fig. S4). Biomass formation at 10 h decreased with increased vial filling (Fig. 3), the optical density dropping from 1.10 ± 0.01 at a filling ratio of 24% to 0.50 ± 0.01 at 96% (Fig. 3A). This observation is consistent with the well-established dependence of *E. coli* growth on oxygen availability, where aerobic respiration supports higher ATP yields and high biomass accumulation, whereas oxygen limitation reduces growth and promotes fermentative metabolism ^34^. As oxygen availability decreased with increasing vial filling, both the WT and Pdu-expressing strains converted glucose into fermentation products, including ethanol, acetate, and formate (Fig. S5 & S6). These results demonstrate that *pdu* expression is not required for their formation and support their origin in the host’s native fermentative metabolism.

**Figure 3:**
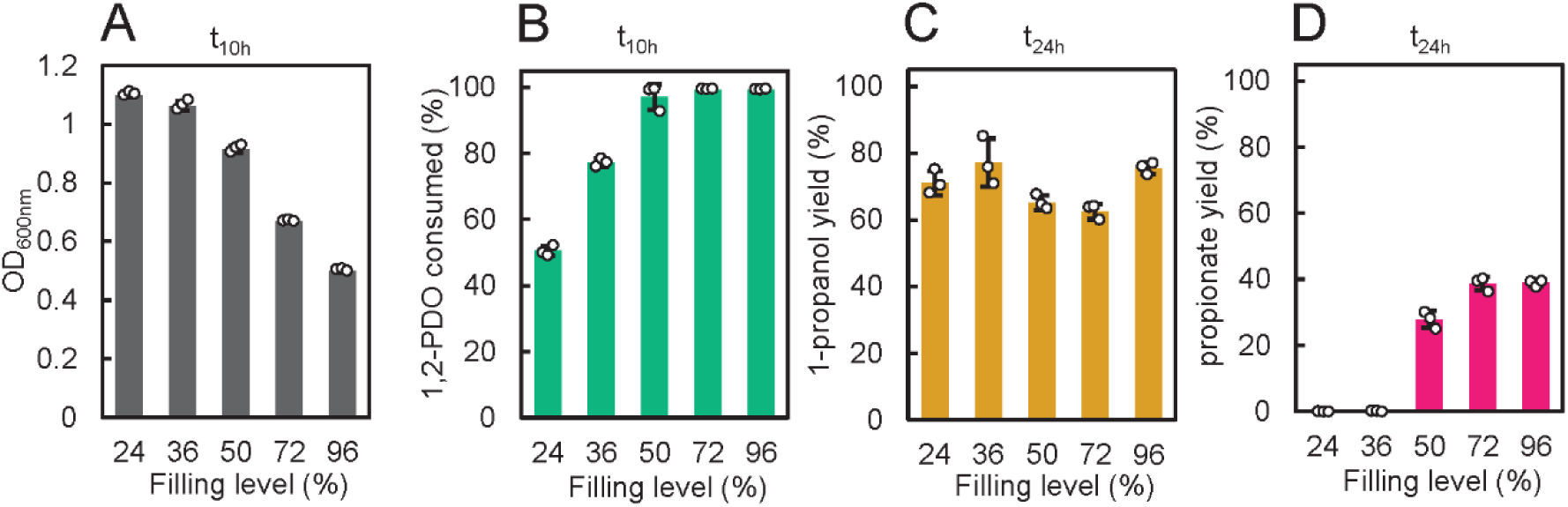
Heterologous Pdu BMCs are more active under microaerobic conditions *in cellula*. *E*. *coli* K-12 BW25113 Pdu was grown in crimp-sealed vials filled with increasing volumes of M9 minimal medium containing D-glucose, 1,2-PDO, cyanocobalamin and tetracycline. Samples were collected over time (Fig. S4). The results obtained after 10 hours of cultivation are shown for (A) optical density at 600 nm (OD₆₀₀nm) and (B) the percentage of 1,2-PDO consumed, determined by ¹H-NMR analysis of the culture medium. The percentages of 1,2-PDO converted into (C) 1-propanol and (D) propionate after 24 h of cultivation are also shown. Data are the averages of n = 3 biological replicates, error bars show standard deviations. Individual values for each biological replicate are overlaid on the corresponding bar graphs.

In contrast, 1,2-PDO utilization was observed only in BW Pdu (Fig. S4 & S7) and appeared strongly enhanced under oxygen-limiting conditions. At 24% filling, only ∼50% of the 1,2-PDO was consumed after 10 h, whereas complete consumption was observed at 50% filling and above (Fig. 3B). Transient accumulation of propionaldehyde in the extracellular medium was detected in all cases (Fig. S4 & S8; see also Supplementary Materials and Methods), with higher concentrations under more oxygen-limited conditions. For example, a maximum 3.65 ± 0.04 mM propionaldehyde was observed after 6 h at a filling ratio of 96%, compared with just 0.86 ± 0.03 mM after 10 h at a filling ratio of 36%. Similar propionaldehyde leakage has been observed in *S. enterica* expressing native Pdu BMCs ^35^. Despite this leakage, carbon recovery from 1,2-PDO in extracellular propionate and 1-propanol remained high (73–100%), indicating limited net loss of intermediates. Interestingly, the proportion of 1,2-PDO converted to 1-propanol remained relatively constant across all conditions (∼60%, Fig. 3C), consistent with observations under fully aerated conditions (Fig. 2E). In contrast, apparent propionate yields increased with increasing vial filling (Fig. 3D). This trend is consistent with reduced propionate reassimilation as oxygen availability decreases. *E. coli* can catabolize propionate through the methylcitrate cycle ^13,36^, and propionate utilization has been demonstrated under oxic conditions but is restricted under anoxic conditions^37^. At lower filling volumes, greater oxygen availability may therefore favor the reassimilation of propionate produced through the Pdu pathway, reducing its net extracellular accumulation. At higher filling volumes, oxygen limitation may reduce reassimilation, leading to greater extracellular accumulation. However, the involvement of the methylcitrate cycle in propionate assimilation remains to be verified under our experimental conditions.

Overall, these results demonstrate that oxygen availability strongly influences Pdu BMC activity, with enhanced 1,2-PDO utilization under microaerobic conditions. This observation supports the notion that key enzymes within the Pdu pathway are oxygen-sensitive even *in cellula*. Subsequent experiments were performed with a 50% filling ratio, which provided an optimal balance between biomass formation and metabolic activity.

### Heterologous Pdu BMCs are metabolically versatile

Pdu BMC’s ability to process a range of diols and triols remains putative, with little direct experimental evidence ^23–25^. To systematically evaluate substrate scope, we tested eleven compounds with different carbon chain lengths and functionalization: five linear 1,2-diols (1,2-PDO as a control (C3), 1, 2-ethanediol, also called ethylene glycol (EG, C2), 1,2-butanediol (1,2-BDO, C4), 1,2-pentanediol (1,2-PTDO, C5) and 1,2-hexanediol (1,2-HDO, C6), three modified diols (2,3-butanediol (2,3-BDO, C4), 3-amino-1,2propanediol (1,2-PDO-A, C3) and 3-chloro-1,2-propanediol (1,2-PDO-Cl, C3); and three triols (glycerol, 1,2,4-butanetriol (1,2,4-BTO, C4) and 1,2,6-hexanetriol (1,2,6-HTO, C6). BW WT and BW Pdu strains were grown under the previously established microaerobic conditions in M9 medium supplemented with D-glucose, cyanocobalamin and the test compounds. Tetracycline was included at inoculation to induce *pdu* expression.

NMR analysis of culture supernatants revealed that five of the eleven substrates were consumed by BW Pdu (Fig. S9 & S10): 1,2-PDO, EG, 1,2-BDO, 2,3-BDO and glycerol, albeit with markedly different efficiencies. BW WT showed only marginal consumption of 1,2-BDO and no detectable consumption of the other three diols (1,2-PDO, EG and 2,3-BDO) (Fig. S11), confirming that their metabolism in BW Pdu is largely dependent on the heterologous Pdu system. The remaining six substrates (1,2-PTDO, 1,2-HDO, 1,2-PDO-A, 1,2-PDO-Cl, 1,2,4-BTO and 1,2,6-HTO) were not tested in BW WT because they were not consumed by BW Pdu (Fig. S12). Substrate uptake rates were estimated in BW Pdu using PhysioFit ^38^ and are summarized in Table 2. Among the diols, 1,2-PDO was the most rapidly consumed (qS = −6.98 ± 0.61 mmol gDW⁻¹·h⁻¹), followed by EG (−4.06 ± 0.34 mmol gCDW⁻¹·h⁻¹) and 1,2-BDO (−0.81 ± 0,21 mmol gCDW⁻¹·h⁻¹), while 2,3-BDO was only weakly utilized (qS too low to be determined).

**Table 2:** Extracellular fluxes of strains producing Pdu BMCs. All extracellular fluxes were calculated using Physiofit ^38^. Percentage conversion was calculated from the measured fluxes. When fluxes could not be determined (Not applicable; NA), conversion was calculated from the fraction of substrate converted to products at the end of the experiment. In some cases, the aldehyde was not detected (ND). The extracellular fluxes are expressed in mmol·gCDW⁻¹·h⁻¹, corresponding to millimoles of substrate consumed or product formed per gram of cell dry weight per hour. By convention, negative values indicate substrate consumption, whereas positive values indicate production. Related charts are shown in Fig. S9, S12, S16 and S17.

| **Strain** | **Subs-trate** | **Extracellular flux in mmol/gCDW/h** |  |  |  | **Conversion in %** |  |  |
|---|---|---|---|---|---|---|---|---|
|  |  | **Substrate** | **Alcohol** | **Acid** | **Aldhehyde** | **Alcohol** | **Acid** | **Aldhehyde** |
| BW Pdu | 1,2-PDO | - 6.98 ± 0.61 | 3.58 ± 0.34 | 1.53 ± 0.24 | 0.76 ± 0.11 | 51 ± 2 | 22 ± 1 | 11 ± 1 |
|  | EG | - 4.06 ± 0.34 | 2.21 ± 0.32 | 0.03 ± 0.12 | 0.85 ± 0.19 | 55 ± 9 | 1 ± 10 | 21 ± 11 |
|  | 1,2-BDO | - 0.79 ± 0.11 | 0.10 ± 0.11 | 0.34 ± 0.10 | ND | 13 ± 2 | 44 ± 17 | ND |
|  | 2,3-BDO | NA | NA | NA | ND | Butanone: 53 ± 7 |  | ND |
|  | Glycerol | - 9.47 ± 0.85 | 1.98 ± 0.36 | 0.22 ± 0.04 | 1.25 ± 0.15 | 21 ± 2 | 2 ± 1 | 13 ± 1 |
| BW Gly<sup>-</sup> Pdu | Glycerol | - 1.30 ± 0.20 | 0.59 ± 0.12 | 0.50 ± 0.11 | 0.00 ± 0.03 | 46 ± 6 | 39 ± 2 | 0 ± 5 |
| BW Pdu* | 1,2-PDO | - 1.11 ± 0.15 | 0.26 ± 0.04 | 0.69 ± 0.07 | ND | 24 ± 2 | 63 ± 9 | ND |
|  | 1,2-BDO | NA | NA | NA | ND | 25 ± 3 | 49 ± 8 | ND |
|  | 1,2-PTDO | NA | NA | NA | ND | 15 ± 1 | 48 ± 1 | ND |
| BW Pdu Δ*pduPQ* | 1,2-PDO | - 6.59 ± 0.48 | 3.93 ± 0.61 | 0.27 ± 0.05 | 3.98 ± 0.61 | 61 ± 17 | 4 ± 1 | 61 ± 14 |
|  | EG | - 1.55 ± 0.28 | 1.07 ± 0.16 | - 1.83 ± 0.07 | 0.77 ± 0.19 | 69 ± 5 | - 119 ± 14 | 50 ± 4 |
|  | 1,2-BDO | - 0.39 ±0.04 | 0.06 ± 0.01 | 0.29 ± 0.02 | ND | 15 ± 5 | 74 ± 6 | ND |
| BW Gly<sup>-</sup> Pdu Δ*pduPQ* | Glycerol | - 0.40 ± 0.06 | 0.19 ± 0.09 | 0.04 ±0.02 | 0.07 ± 0.06 | 50 ± 10 | 12 ± 6 | 16 ± 10 |

Consistent with the canonical pathway, the metabolized 1,2-diols were converted into a mixture of corresponding alcohols and carboxylic acids via aldehyde intermediates (Table 2). NMR-based quantification (Fig. S9) showed that 1,2-PDO yielded 1-propanol (51% of carbon) and propionate (22%), EG yielded ethanol (55%) and acetate (1%), and 1,2-BDO yielded butyrate (39%) and 1-butanol (13%). Aldehydes were detected in the extracellular medium for all tested substrates except 1,2-BDO (Table 2). None of these products were observed in BW WT cultures (Fig. S11). While product distributions varied between substrates, these results demonstrate that Pdu BMCs support promiscuous metabolism with multiple 1,2-diols. Additionally, butanone was detected during cultivation with 2,3-BDO (Fig. S9D), consistent with PduCDE-mediated dehydration, as previously reported for related diol dehydratases^39^. Its accumulation suggests limited downstream conversion. This is consistent with the established function of PduP as a CoA-acylating aldehyde dehydrogenase^40^, since butanone is a ketone and is therefore not expected to undergo the same reaction.

Among the tested triols, only glycerol was consumed by BW Pdu (qS = −9.47 ± 0.85 mmol gCDW⁻¹·h⁻¹) (Table 2, Fig. S10B), with concomitant production of 3-HP (2% of the carbon from glycerol) and 1,3-PDO (21%). 3-HPA was also detected (13%). Native *E*. *coli* pathways also contributed to glycerol metabolism ^41^, as reflected by glycerol consumption in BW WT, albeit at a lower rate (qS = −3.70 ± 0.69 mmol gCDW⁻¹·h⁻¹), and without production of 3-HPA, 3-HP or 1,3-PDO (Fig. S10A). This indicates a significant contribution of Pdu BMCs to glycerol metabolism in BW Pdu. To further isolate the role of the Pdu system from central glycerol metabolism, we integrated the P*_tet_::pdu* construct into a glycerol-negative background (BW Δ*dhaKLM,* Δ*glpk* and Δ*gldA*; see Supplementary Materials and Methods), generating BW Gly^−^ Pdu (Table 1). As expected, the BW Gly^−^ control strain did not consume any glycerol (Fig. S10C), while in BW Gly^−^ Pdu, glycerol utilization was partially restored, but at a 7.2-fold lower rate (Table 2, Fig. S10D) (−1.30 ± 0.20 mmol gCDW⁻¹ h⁻¹) than in BW Pdu. Notably, carbon flux toward 3-HP (39% of the carbon from glycerol) and 1,3-PDO (46%) were higher than in BW Pdu, although absolute titers were lower (Table 2, Fig. S10D). The reduced uptake rate in the Gly⁻ background may reflect impaired expression of the glycerol transporter GlpF, which is regulated by the repressor GlpR, whose inhibition is reduced upon binding glycerol-3-phosphate, thereby activating transcription of the *glp* regulon ^42^. Glycerol kinase (GlpK) is absent in the Gly^−^ strains and hence no glycerol-3-phosphates accumulates, potentially limiting GlpF expression.

Taken together, our findings show that heterologous Pdu BMCs have substantial metabolic promiscuity. EG was the most efficiently utilized of the tested substrates, other than 1,2-PDO, while glycerol metabolism was influenced by host regulatory constraints. For 1,2-PDO, EG, and glycerol, product formation was consistently biased toward alcohol production, whereas 1,2-BDO conversion was less efficient and promoted acid production. For 2,3-BDO in contrast, the pathway remained incomplete, highlighting substrate-dependent limitations in downstream enzymatic steps.

### Pore permeability does not prevent entry of longer diols

We next investigated why six of the eleven tested compounds were not metabolized by BW Pdu. Substrate utilization can be restricted at multiple stages, including by transport across the cellular membrane, passage through the BMC shell and access to encapsulated Pdu enzymes. Previous work has shown that a mutated form of the *L*. *reuteri* PduCDE diol dehydratase, when expressed in the cytosol of *E*. *coli*, can convert several exogenous substrates *in cellula* (i.e. EG, glycerol, 1,2-BDO, 1,2-PTDO and 1,2-HDO) ^26^. This suggests that at least some of the compounds not metabolized by BW Pdu (i.e. 1,2-PTO and 1,2-HDO) can reach the cytosol. We therefore evaluated whether diffusion across the BMC shell pores could represent a limiting step.

Since direct measurements of metabolite diffusion across BMC shells are not yet feasible, we used molecular dynamics simulations to estimate BMC permeability by tracking permeation events of small molecules across a model sheet of BMC shell proteins. This sheet consisted of a PduA homohexamer, a PduJ homohexamer, a PduA/J heterohexamer and a PduB’ trimer (Fig. 4A & 4B), approximating a simplified facet of the Pdu BMC polyhedral shell ^33^. Simulations were performed for seven compounds: EG, glycerol, 1,2-PDO (Fig. S13), 1,2-BDO, 1,2-PTDO, 1,2-HDO and 1,2,6-HTO. As expected, the substrates crossed the shell through pores located at or close to the center of the protein tiles (Fig. 4C). Transition events, defined as any event in which a small molecule diffuses from one side of the protein sheet to the other through a tile pore, were measured and recorded for all simulations. Between 13 and 603 transition events through individual shell tiles were observed per metabolite over 6 μs of sampling (Table 3). The permeability coefficients were determined based on the transit rate, normalized by the compound concentration (see Materials and Methods and Supplementary Materials and Methods). The range of permeability coefficients was surprisingly small (Table 3), with no more than a 50-fold difference between the fastest and slowest combinations of protein component and metabolite.

**Figure 4:**
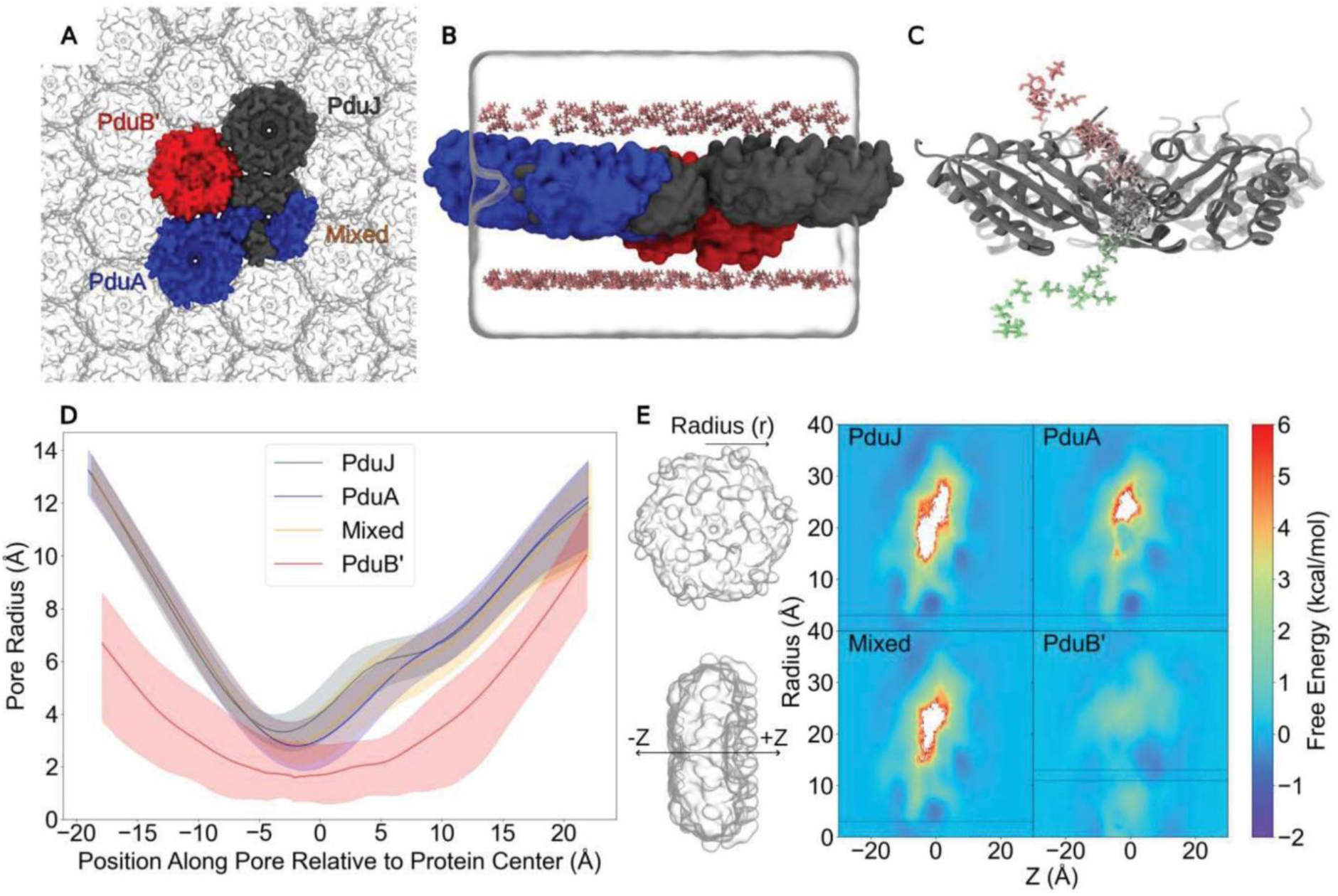
Analysis of Pdu BMC shell permeability by molecular dynamics simulations. (A) Top view of the simulated *C*. *freundii* Pdu BMC sheet consisting of a PduA homohexamer (blue), a PduJ homohexamer (grey), a mixed PduA/PduJ heterohexamer (blue and grey) and a PduB’ homotrimer (red) and (B) side view of a single periodic system with 1,2-PDO colored in pink. (C) Example of a 1,2-PDO transit event. This timeseries shows a single 1,2-PDO molecule crossing through the central pore of the PduJ homohexamer, with pinker colors indicating the start of the trajectory and greener colors indicating the end. (D) Distribution of pore radii throughout the simulations investigating the permeability of 1,2-PDO. The dark line at the center represents the mean radii while the shaded region around this line represents the 5^th^ and 95^th^ percentile radii. Proteins were centered by mass at Z = 0.A single representative radius was visualized for the three pores of PduB’ to avoid redundancy. (E) Free energy profiles for 1,2-PDO transiting through the pore in two dimensions. Free energies were determined by inverting the probability distributions of metabolite positions and are corrected for increasing protein surface areas at larger radii. Regions colored in white represent locations where sampling did not occur and where the probability distribution was thus undefined. Negative free energy values at large radii are likely due to the imperfect packing of proteins in the tiled sheet, which is a non-native assembly.The pores for the hexameric proteins, PduJ, PduA, and Mixed, are located at r = 0 Å, while Pdub’, the trimeric protein, has 3 small pores located at r = 12 Å, and their boundaries are indicated by dashed lines.

**Table 3:** Permeability values for selected metabolites across Pdu BMC shell protein oligomers. Molecular dynamics simulations were performed separately for each metabolite. The number of transitions across individual shell oligomers of the reconstructed sheet (Fig.4, B) was counted over 1 µs of simulation. The analysis involved six independent replicates, i.e. 6 µs of simulation in total. Permeability coefficients (cm·s^−1^) were calculated as detailed in the Materials and Methods section. Values (± standard deviations) are the averages of n = 6 replicates with errors indicating SD.

| Metabolite | Tile | Transitions | P |
|---|---|---|---|
| 1,2-PDO | PduA | 200 | 0.13 ± 0.02 |
|  | PduJ | 414 | 0.27 ± 0.02 |
|  | Mixed | 280 | 0.18 ± 0.02 |
|  | PduB' | 154 | 0.10 ± 0.04 |
| EG | PduA | 361 | 0.24 ± 0.04 |
|  | PduJ | 603 | 0.40 ± 0.03 |
|  | Mixed | 576 | 0.38 ± 0.06 |
|  | PduB' | 192 | 0.13 ± 0.07 |
| 1,2-BDO | PduA | 113 | 0.07 ± 0.01 |
|  | PduJ | 250 | 0.16 ± 0.01 |
|  | Mixed | 287 | 0.19 ± 0.03 |
|  | PduB' | 34 | 0.02 ± 0.01 |
| 1,2-PTDO | PduA | 91 | 0.06 ± 0.01 |
|  | PduJ | 183 | 0.12 ± 0.02 |
|  | Mixed | 154 | 0.10 ± 0.02 |
|  | PduB' | 16 | 0.01 ± 0.01 |
| 1,2-HDO | PduA | 51 | 0.03 ± 0.01 |
|  | PduJ | 122 | 0.08 ± 0.01 |
|  | Mixed | 156 | 0.10 ± 0.01 |
|  | PduB' | 35 | 0.02 ± 0.01 |
| Glycerol | PduA | 201 | 0.13 ± 0.02 |
|  | PduJ | 404 | 0.27 ± 0.02 |
|  | Mixed | 374 | 0.13 ± 0.02 |
|  | PduB' | 70 | 0.05 ± 0.02 |
| 1,2,6-HTO | PduA | 55 | 0.04 ± 0.01 |
|  | PduJ | 140 | 0.09 ± 0.01 |
|  | Mixed | 133 | 0.09 ± 0.02 |
|  | PduB' | 13 | 0.01 ± 0.00 |

The PduJ homohexamers and PduA/PduJ mixed heterohexamers were generally more permeable than the PduA homohexamers, which were themselves far more permeable than the PduB’ trimers. As we track permeation events, compounds permeate through the central pore in the hexameric tiles, while we find that permeation events are localized away from the center in the PduB’ trimers. We compare the pore radii along the paths that permeants follow using the HOLE ^43^ program (Fig. 4D). The small pore for PduB’ is off axis and within a single trimeric unit, with a closed central pore rather than a single large central open pore as in other BMC shell protein trimers ^44^. A similarly “closed” conformation appears in the crystal structure of *Lactobacillus reuteri* PduB ^18^. While PduB and PduB’ may also adopt an “open” conformation, similar to EutL’s in the Eut BMC system ^19^, the conditions required to trigger this structural shift remain unknown. Subsequent analyses were therefore performed on the “closed” conformation of PduB’. By tracking individual particles during the simulation, we can map probabilities for small molecules compared to their position along an aligned protein structure. Free energy profiles can be determined from these probabilities, highlighting again that the lowest free energy path through the PduB’ trimer was along the three small pores approximately 12 Å from the protein center (region bounded by dotted lines in Fig. 4E). HOLE analysis (Fig. 4D) and the inverted probability distributions for 1,2-PDO across the shell (Fig. 4E) both showed that the PduJ, PduA and the mixed hexamer tiles had larger central pore radii, allowing metabolite flux through the reconstructed Pdu BMC shell protein sheet through the hexamers ^15,45^. Indeed, there is little barrier for permeation up to 2.5 Å from the pore center in the hexamer PMFs.

These lower energy paths along the 2-D PMFs allow us to construct and analyze 1-D PMFs at the respective pore locations (Fig. S14). PduJ and the mixed hexamer share a similar free energy profile and greater permeability as compared to PduA which has a markedly less favorable PMF. Notably, the PMF for PduB’ shows a favorable pathway for diffusion with only moderate energy barriers, seemingly at odds with the reported permeability coefficients. However, further analysis on mean transit times (Fig. S15), where PduB’ transit can take tens of nanoseconds, suggests that the greater constriction offered by the smaller pore results in an increase in interactions with the protein structure, delaying permeation events. Indeed, diffusion through PduB’ accounts for almost all the transition events lasting more than 10 ns, with some outliers reaching up to 40 ns. These long transition events likely occlude the pores from further permeation and reduce the overall permeability coefficients of PduB’.

Importantly, the calculated permeability coefficients were relatively high for all tested substrates (Table 3), indicating that longer chain diols and triols are not blocked from entering the Pdu BMC. For example, permeability through the central pore of the PduJ homohexamer was estimated at 0.12 and 0.09 cm·s^−1^ for 1,2-PTDO and 1,2-HDO, respectively, approximately one third the value obtained for the native substrate, 1,2-PDO (0.27 cm·s^−1^), but far from negligible. These values would not support the formation of large concentration gradients across the shell during substrate consumption, and altogether, these results suggest that shell permeability is unlikely to be the primary limiting factor for the utilization of longer-chain diols such as 1,2-PTDO and 1,2-HDO within Pdu BMCs.

### Heterologous Pdu BMCs can be redesigned to act on longer chain diols

Since our molecular dynamics analysis indicates that 1,2-PTDO and 1,2-HDO can likely enter Pdu BMCs, their utilization must be restricted in downstream enzymatic conversion. Indeed, when expressed recombinantly in the cytosol of *E*. *coli*, the *L*. *reuteri* PduCDE complex does not metabolize diols longer than 1,2-BDO ^26^. However, two point mutations in PduC (S302A and Q337A) have been shown to enhance activity toward longer-chain diols of up to six carbons ^26,46^. We therefore introduced the equivalent mutations (S301A Q336A) into the *C. freundii* PduC, generating the BW Pdu* strain (Table 1). BW Pdu* strain was grown under the previously established microaerobic conditions in M9 medium supplemented with D-glucose, cyanocobalamin and the test compounds. Tetracycline was included at inoculation to induce *pdu* expression.

As shown in Table 2 and Fig. S16, and consistent with previous reports ^26,46^, these mutations markedly reduced 1,2-PDO consumption, with a ∼6-fold lower uptake rate in BW Pdu* (qS = −1.11 ± 0.15 mmol gCDW⁻¹·h⁻¹) than in BW Pdu (qS = −6.98 ± 0.61 mmol gDW⁻¹·h⁻¹). Product distribution was also altered, with 24% of the recovered 1,2-PDO carbon directed toward 1-propanol and 63% toward propionate. Propionaldehyde was not detected in the culture supernatants. For 1,2-BDO, BW Pdu* behaved similarly to BW Pdu (Fig. S9C & S16B), consuming more than 70% of the substrate within 48 h and favoring butyrate (49%) over 1-butanol (24%). In contrast, BW Pdu* partially metabolized 1,2-PTDO, of which roughly 50% was consumed after 48 h, yielding 1-pentanol (15% of carbon) and valeric acid (48%) (Table 2, Fig. S16C), whereas no detectable utilization of this substrate was observed with BW Pdu (Fig. S12A). These results confirm that 1,2-PTDO can cross both the cellular membrane and the Pdu BMC shell, and that its lack of utilization is due to enzymatic constraints. In contrast, no significant uptake of 1,2-HDO was observed in BW Pdu* (Fig.S16D), although trace amounts of caproic acid were detected. This suggests that additional limitations, potentially including reduced transport across cellular or BMC membranes, lower enzyme affinity, or slower catalytic turnover, restrict the metabolism of longer-chain diols.

Although targeted mutations in PduCDE reduced the conversion rate of the native substrate, 1,2-propanediol, they extended the substrate range of the Pdu BMC system to one additional substrate, whose aldehyde intermediate could be further processed by the downstream enzymes. This finding suggests that downstream enzyme promiscuity can accommodate changes in substrate specificity at the initial catalytic step, albeit with a trade-off in native substrate conversion.

### Contribution of cytosolic enzymes to aldehyde fate

Aldehydes were detected in BW Pdu culture supernatants for nearly all metabolized polyols (Fig. S9 & S10), raising questions about their metabolic fate and the potential contribution of host cytosolic enzymes to their conversion. We therefore investigated the extent to which endogenous activities could compensate for the loss of aldehyde-processing reactions within Pdu BMCs. To this end, we constructed a BW Pdu Δ*pduPQ* mutant (Table 1, Fig. S2B & S2D) and characterized its metabolic activity under the previously established microaerobic conditions in M9 medium supplemented with D-glucose, cyanocobalamin and the test compounds. Tetracycline was included at inoculation to induce *pdu* expression.

BW Pdu Δ*pduPQ* assimilated 1,2-PDO (qS = −6.59 ± 0.48 mmol gDW⁻¹·h⁻¹) at a slightly lower rate than BW Pdu (Table 2, Fig. S17A). Strikingly, propionaldehyde excretion (qP = 3.98 ± 0.61 mmol gDW⁻¹·h⁻¹) increased 5.2-fold, indicating that endogenous enzymes could not fully compensate for the loss of PduP and PduQ. Nevertheless, substantial 1-propanol production (qP = 3.93 ± 0.61 mmol gDW⁻¹·h⁻¹), together with modest propionate formation, suggests that propionaldehyde can still be reduced by cytosolic enzymes, whereas its oxidation is severely limited in the absence of PduP. The mutant assimilated EG (qS = -1.55 ± 0.28 mmol gDW⁻¹·h⁻¹) 2.6-fold more slowly than BW Pdu, while excreting acetaldehyde at a comparable rate (qP = 0.77 ± 0.19 mmol gDW⁻¹·h⁻¹) (Table 2, Fig. S17B). The reduced EG conversion rate, together with the persistent acetaldehyde accumulation, suggest that endogenous aldehyde-processing activities could not fully compensate for the deleted BMC-associated enzymes, PduP and PduQ. 1,2-BDO assimilation also decreased in BW Pdu Δ*pduPQ* (qS = -0.38 ± 0.04 mmol gDW⁻¹·h⁻¹) (Table 2, Fig. S17C). No butyraldehyde was detected and butyrate production (qP = 0.29 ± 0.03 mmol gDW⁻¹·h⁻¹) remained similar to that of BW Pdu, whereas 1-butanol production decreased markedly. These results suggest that endogenous activities could compensate for the loss of PduP-mediated butyraldehyde oxidation, but provided limited compensation for PduQ-mediated reduction. Finally, we assessed the impact of the *pduPQ* deletion in the BW Gly⁻ background (Tables 1 and 2, Fig. S17D). BW Gly^-^ Pdu Δ*pduPQ* consumed glycerol at a lower rate (qS = -0.40 ± 0.06 mmol gDW⁻¹·h⁻¹) than BW Gly^-^ Pdu. The main product was 1,3-PDO (qP = 0.19 ± 0.09 mmol gDW⁻¹·h⁻¹), while only small amounts of 3-HP were detected. Moreover, 3-HPA excretion (qP = 0.07 ± 0.06 mmol gDW⁻¹·h⁻¹) increased compared with the parental strain. This profile indicates that endogenous enzymes can reduce 3-HPA to 1,3-PDO, whereas its oxidation to 3-HP is severely limited.

*E. coli* possesses several aldehyde reductases and dehydrogenases with broad, overlapping substrate specificities, providing potential alternative routes for processing the aldehydes generated by PduCDE. On the reductive side, YahK, YjgB and YqhD have been shown to catalyze the NADPH-dependent reduction of acetaldehyde, propionaldehyde and butyraldehyde^47^. On the oxidative side, AldB exhibits NADP⁺-dependent activity towards acetaldehyde and propionaldehyde^48^, while AldH displays broad substrate specificity that includes 3-HPA, propionaldehyde and butyraldehyde ^49^. Furthermore, deleting *yqhD* markedly affected 3-HPA conversion to 1,3-PDO in *E*. *coli* ^50^, supporting a significant contribution of endogenous YqhD to 3-HPA conversion.

Overall, our results reveal substrate-dependent differences in the capacity of cytosolic enzymes to process polyol-derived aldehydes. Previous studies identified several endogenous enzymes with relevant reductive or oxidative activities *in vitro* and, in one case, *in cellula*. Although their individual contributions under our experimental conditions remains to be established, our results demonstrate that these endogenous activities cannot fully compensate for the loss of PduP and PduQ. This finding highlights the importance of downstream aldehyde-processing reactions for efficient polyol conversion by the intact Pdu BMC.

### Factors influencing product distribution in BW Pdu

Our results indicate that, regardless of the substrate, product distribution after polyol conversion is shifted rather than balanced – either toward the alcohol or the corresponding carboxylic acid (Table 2). One possible explanation lies in the interplay between aldehyde generation by PduCDE, downstream processing within the BMCs and conversion by host cytosolic enzymes. When aldehyde generation exceeds the capacity of the downstream BMC-associated enzymes, intermediates may accumulate and escape into the cytosol, where endogenous activities can further influence the final alcohol- to-acid ratio. The product profiles of BW Pdu Δ*pduPQ* suggest that endogenous activities compensate more effectively for the reduction of propionaldehyde, acetaldehyde and 3-HPA than for their oxidation (Fig. S17). Leakage of these intermediates could therefore favor alcohol formation. In contrast, endogenous processing of butyraldehyde appears to favor oxidation, potentially increasing the butyrate fraction. Consistent with this interpretation, rapid 1,2-PDO consumption by BW Pdu was associated with detectable propionaldehyde accumulation and an alcohol-favored product distribution (Table 2, Fig. S4). By comparison, BW Pdu*, which consumed 1,2-PDO more slowly, showed no detectable extracellular propionaldehyde and a shift towards acid production (Table 2, Fig. S11). Similarly, slower glycerol consumption by BW Gly⁻ Pdu than by BW Pdu was accompanied by lower aldehyde accumulation and a greater acid fraction. The relatively slow conversion of 1,2-BDO by both strains also favored butyrate over 1-butanol formation (Table 2), consistent with the apparent oxidative capacity of host enzymes towards butyraldehyde.

Product distributions may also be influenced by the consumption of Pdu pathway-derived acids and alcohols through endogenous *E. coli* metabolic activities. For instance, propionate can enter central metabolism via the methylcitrate cycle ^13,36^ and acetate can be metabolized through the Pta–AckA and the Acs pathways^51^. During EG conversion, acetate utilization within cytosol could limit its excretion, as observed during Eut BMC metabolism^52^. Alcohols may also undergo secondary conversion. For instance, oxidation of 1-butanol to butyraldehyde by endogenous alcohol dehydrogenases such as AdhE^53^, followed by oxidation to butyrate by an aldehyde dehydrogenase such as AldH^49^, could contribute to butyrate formation. However, this hypothesis remains to be tested under our experimental condition. Finally, BW Gly⁻ and BW Gly⁻ Pdu were cultivated with ^13^C-labeled glycerol under the same conditions (see Supplementary Materials and Methods). Although both 1,3-PDO and 3-HP were labelled, no 13C incorporation was detected in cellular amino acids (Fig. S18). This result demonstrates that 3-HP was not used by anabolism.

Together, these observations suggest that final product distributions reflect both the balance of catalytic activities within Pdu BMCs and their interactions with host metabolism, including aldehyde conversion and potential product reutilization.

## Conclusion

The metabolic promiscuity observed in this study extends beyond the relaxed specificity of individual enzymes and emerges at the level of the entire encapsulated pathway. Although substrate promiscuity has been described for isolated enzymes, including PduCDE and related aldehyde-processing enzymes ^26^, direct experimental evidence at the level of functional BMCs is lacking. Our results demonstrate that non-native substrates can be accepted, transported across the shell, and sequentially metabolized by the complete Pdu BMC system. This coordinated flexibility reveals a higher-order “pathway promiscuity”, in which substrate scope is determined collectively by shell permeability and the catalytic plasticity of multiple encapsulated enzymes. To our knowledge, these findings are the first experimental demonstrations of pathway-level promiscuity in an intact BMC, casting the Pdu BMC as a metabolically adaptable module rather than just a single-substrate pathway. Whether this promiscuity provides a selective advantage *in vivo* remains to be established, but it may enhance the capacity of BMCs to adapt to fluctuating nutrient environments and thereby confer metabolic robustness. From a biotechnological perspective, this intrinsic flexibility broadens the repertoire of substrates amenable to BMC-based bioconversion and provides a basis for further expansion through enzyme engineering, as demonstrated here. Such versatility offers opportunities for the production of value-added chemicals and biological upcycling. For example, 1,3-PDO and 3-HP are industrial platform chemicals with applications in polymer production and chemical manufacturing, whereas PET-derived ethylene glycol can serve as a substrate for plastic waste valorization ^54,55^.

Nevertheless, our results show that this flexibility operates in crosstalk rather than independently from the host cell. Metabolites, reactive intermediates, and possibly cofactors can cross the semi-permeable BMC shell, creating dynamic exchanges between the compartment and the cytosol. In the Pdu BMC expressed in *E. coli*, this imperfect orthogonality manifests in the activity of native cytosolic enzymes (e.g. responsible for aldehyde detoxification and products consumption), which can further transform metabolites released from the compartment. Likewise, although Pdu BMCs encapsulate oxygen-sensitive enzymes and are thought to create a locally microaerobic environment, our data indicate that oxygen negatively affects pathway performance *in cellula*, suggesting that it can partially access the compartment interior.

Importantly, product distribution is shaped not only by metabolite leakage to the cytosol but also by the substrate-dependent kinetic properties of the encapsulated enzymes themselves. High substrate influx may saturate internal enzymatic steps, promoting accumulation and escape of aldehyde intermediates, while the relative formation of alcohol and acid products depends on the balance between competing enzymatic conversion rates inside the BMC. Consequently, the final metabolic output depends on the interplay between substrate transport, internal pathway kinetics, intermediate leakage, and host metabolism, rather than from the internal stoichiometry of the compartment alone.

Fully exploiting Pdu BMCs as programmable biotechnological modules will require careful control of substrate influx, host metabolic backgrounds, and of the catalytic properties of encapsulated enzymes. In this context, *in vitro* reconstitution systems represent a promising complementary strategy to dissect and optimize these parameters under controlled conditions. By decoupling BMC function from native cellular metabolism, such systems could enable detailed analyses of pathway flux, cofactor exchange, and shell permeability, guiding the rational engineering of next-generation BMCs.

## Material and Methods

### Bacterial strains and cultivation conditions

All bacterial strains employed in this study are listed in Table 1. Cells were cultivated either in LB medium ^56^ or in M9 minimal medium ^52^. The M9 medium was supplemented with thiamine hydrochloride (100 mg/L), cyanocobalamin (200 nM), tetracycline (3 µg/L), and D-glucose (5 mM) as the carbon and energy source. Ethylene glycol, 1,2-propanediol, glycerol, 1,2-butanediol, 2,3-butanediol, 1,2-pentanediol, 1,2-hexanediol, 3-amino-1,2-propanediol, 3-chloro-1,2-propanediol, 1,2,4-butanetriol or 1,2,6-hexanetriol were added to the final concentration of 10 mM. All chemicals were obtained from Sigma-Aldrich (France). All media components were sterilized by autoclaving except thiamine hydrochloride, trace elements, cyanocobalamin, tetracycline, glucose, and diols/triols, which were filter sterilized (0.2 µm pore size; Minisart; Sartorius).

Glycerol stocks stored at −70°C were streaked onto LB agar plates to obtain isolated colonies. Single colonies were used to inoculate LB precultures grown at 37°C with double orbital shaking (Innova 4230 incubator; New Brunswick Scientific; 200 rpm) over the day. Cells from the LB precultures were used to inoculate M9 minimal medium supplemented with glucose and grown overnight at 37°C 200 rpm. These cultures were then used to inoculate fresh M9 medium at an initial OD_600nm_ of 0.05. Cultivations were performed at 37°C with double orbital shaking at 200 rpm under either aerobic condition (baffled flasks) or microaerobic condition (sealed bottles) as indicated in the text. Samples (1 mL) were collected at the indicated time points. Cell growth was monitored by measuring the optical density at 600 nm (OD_600nm_) using a Genesys 150 spectrophotometer (Thermo Fisher Scientific). Samples were centrifuged at 16,500 × g for 30 s at room temperature to separate cells from supernatant. The supernatant was collected and stored at −20°C until further analysis.

### Preparation of competent cells and transformation

Bacterial strains were transformed by chemical transformation using the rubidium chloride method ^57^. For transformation, 100 ng of plasmid DNA was added to 50 µL of competent cells and incubated on ice for 20–30 min. Cells were heat shocked at 42°C for 90 s and then returned to ice for 2 min. LB medium (200 µL) was added, and cells were allowed to recover for 45–60 min (or 120 min for strains carrying two plasmids) at 37°C with shaking at 200 rpm (or at 30°C when strains carrying the thermosensitive pCas plasmid were used).

### CRISPR-Cas9 genome editing

Genome editing, including integration of the *pdu* operon, introduction of point mutations in *pduC* and deletion of *pduP* and *pduQ*, was performed following a protocol adapted from ^58^. The *E. coli* BW25113 wild-type strain was co-transformed by heat shock with plasmids pTetPdu (Fig. S1) and pCas, which allows expression of the Cas9 protein and the λ Red recombination system. pCas was a gift from Sheng Yang (Addgene plasmid # 62225; http://n2t.net/addgene:62225; RRID: Addgene_62225). Transformants were selected on LB agar supplemented with ampicillin (100 µg/mL) and kanamycin (50 µg/mL) and incubated at 30°C. A single colony was used to inoculate LB medium supplemented with ampicillin (50 µg/mL) and kanamycin (25 µg/mL) and grown at 30°C 200 rpm shaking. At mid-exponential phase (OD_600nm_ ≈ 0.3), expression of the λ Red recombinase was induced by addition of L-arabinose to a final concentration of 10 mM, followed by incubation for 1 h. Competent cells were then prepared and transformed with plasmid pTarget-N20-1. pTarget-N20-1 was designed to target the *cobU*-*yoeG* chromosomal locus of *E*. *coli* K-12; it was generated by cloning a 20 nucleotides long spacer sequence (TGGCGACTATGCACTAGGGA) into pTargetF using In-Fusion assembly (Takara). pTargetF was a gift from Sheng Yang (Addgene plasmid # 62226; http://n2t.net/addgene:62226; RRID:Addgene_62226). Transformants were selected on LB agar supplemented with kanamycin (50 µg/mL) and spectinomycin (100 µg/mL) and incubated at 30°C. Plasmids were subsequently cured as described previously by ^58^ Point mutations in *pduCDE* were introduced by PCR mutagenesis. The resulting mutant sequence was then integrated into the chromosomal *pdu* operon of BW Pdu using the same CRISPR-Cas9 genome editing system. *pduP* and *pduQ* were deleted from BW Pdu and BW Gly^-^ Pdu with a similar pipeline.

### NMR analysis and fluxes calculation

The culture supernatants were analyzed by 1D proton nuclear magnetic resonance (1D ^1^H-NMR) to quantify substrate consumption and extracellular metabolite production. First, each culture supernatant (180 μL) was mixed with 20 μL of a 10 mM 3-trimethylsilylpropionic-2,2,3,3-d4 sodium salt (TSP-d4; Sigma) solution prepared in D_2_O. TSP-d4 served as an internal standard for quantification and was additionally counter-quantified using a 1 g/L succinate analytical standard. Analyses were performed on a 500 MHz Avance III spectrometer (Bruker, Rheinstetten, Germany) equipped with a 5 mm QPCI cryogenic probe using a zgpr30 sequence with water presaturation before acquisition. The acquisition parameters were as follows: 286 K, 128K points, 8 s relaxation time, 2 dummy scans, 32 scans. Free induction decays (FID) were converted into frequency domain spectra by Fourier transformation. All spectra were processed using the TopSpin software (v. 4.0, Bruker). The spectra were manually phased, baseline corrected automatically and aligned and quantified using TSP. The concentrations of different metabolites were calculated using the formula relating peak area (S), concentration and number of protons assigned to the peak 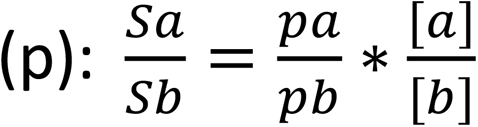, where [*a*] corresponds to the known concentration of TSP-d4 and [*b*] to the concentration of compound of interest b. Substrates uptake and products formation fluxes were calculated with the generated time series data using Physiofit ^38^. All acquired NMR spectra, together with the associated raw data, integration details and quantitative results, are publicly available on ZENODO (https://doi.org/10.5281/zenodo.24468516)

### Molecular dynamics simulations

This part is detailed in supplemental Materials and Methods. In summary, to investigate metabolite permeability through Pdu BMC shell pores, an atomistic model comprising four shell tiles arranged in a planar rhombus configuration was constructed. Tile structures, including PduA, PduJ, a mixed PduA/PduJ heterohexamer, and PduB’, were predicted using AlphaFold 3 based on sequences from *Citrobacter freundii*. Molecular dynamics simulations were performed to evaluate the permeability of glycerol, ethylene glycol, 1,2-propanediol, 1,2-butanediol, 1,2-pentanediol, 1,2-hexanediol, and 1,2,6-hexanetriol. Each metabolite was simulated at 500 mM in six independent 1-µs replicates, yielding 42 µs of cumulative simulation time. Permeability (P) was estimated from the rate (r) of molecular transit events across the shell pores, adjusted for our 500 mM metabolite concentration c. This formulation, where P=r/2c, follows Equation 29^59^.Transit events were identified by tracking metabolite positions along the membrane normal throughout the trajectories. Pore geometries were characterized using the HOLE program and the HoleHelper plugin ^43,44^. To further investigate transport mechanisms, free-energy landscapes were generated from metabolite probability distributions as a function of radial distance from the pore center and position along the pore axis. Together, these simulations provided a quantitative description of metabolite permeability and permeation pathways through Pdu BMC shell pores. The associated molecular simulation data is available on ZENODO (https://doi.org/10.5281/zenodo.19759241).

## Acknowledgements

We thank MetaToul-FluxoMet (Toulouse, France), which is part of the French National Infrastructure for Metabolomics and Fluxomics (https://www.metabohub.fr) and funded by the ANR (MetaboHUB-ANR-11-INBS-0010), for access to MS and NMR facilities. We acknowledge the METi imaging facility (Genotoul-TRI), member of the national infrastructure France-BioImaging supported by the French National Research Agency (ANR-24-INBS-0005 FBI BIOGEN). The authors also thank Edern Cahoreau for his assistance with 2D NMR analyses and data processing, Erwana Harscoet for her support in enzyme engineering, David Haoreau for his help with BMC purification, and Mathias Orlando for his assistance in setting up the cultivation parameters.

The work of DG, LJ, PM, DJ and SH was supported by government funding administered by the National Research Agency under the reference numbers ANR-23-CE44-0038 & ANR-18-EURE-0021. JVV was supported as part of the Center for Catalysis in Biomimetic Confinement, an Energy Frontier Research Center funded by the U.S. Department of Energy (DOE), Office of Science, Basic Energy Sciences (BES), under award DE-SC0023395, and AWS was supported by the National Institutes of Health under award number R35GM155317. Simulations were supported through computational resources and services provided by the Institute for Cyber-Enabled Research at Michigan State University. MW gratefully acknowledges the support of the Biotechnology and Biological Sciences Research Council (BBSRC); this research was funded by the BBSRC Institute Strategic Programme Food Microbiome and Health BB/X011054/1 and project grant BB/Y008456/1.

## Author contribution

D.G., D.J. and S.H. designed and performed the *in cellula* experiments and analyzed the data.

A.W.S. and J.V.V. designed and performed the molecular dynamics analysis and analyzed the data.

L.J. designed and performed the *in vitro* experiments and analyzed the data.

V.S. performed the TEM experiments.

E.D. and M.W. designed and built the plasmid encoded for the pdu BMC operon from *Citrobacter freundii*.

D.G., D.J. P.M. and S.H. wrote the manuscript with contributions from the other authors.

D.J. and S. H. conceptualized the project and designed and supervised the research.

## Data availability

Additional information on the experimental setup as well as detailed results are available from the corresponding author upon request. The raw data supporting the findings presented in the figures and tables are available through Zenodo (https://doi.org/10.5281/zenodo.19759241 & https://doi.org/10.5281/zenodo.24468516). Any strains and plasmids generated during this study are available upon completing a Materials Transfer Agreement.

## Materials & Correspondence

Request should be addressed to stephanie.

