## Supplemental Figures for "Small but Mighty: The Surprising Metabolic Range of a Bacterial Microcompartment"

| Plasmid name | Description | Relevant features | Source |
|---|---|---|---|
| pTetPdu | Chromosomal integration vector | *Pdu* operon from *Citrobacter freundii* under the dependency of an inducible $P_{tet}$ promoter. The operon is flanked by homologous regions for CRISPR/Cas9 integration into the *yoeG/cobU* chromosomal locus of *E. coli* K-12 BW25113. | Kindly provided by Martin Warren |
| pTarget-CobU | gRNA expressing vector | gRNA targeting the *yoeG/cobU E. coli* chromosomal locus for CRISPR/Cas9–mediated genome editing | Kindly provided by Martin Warren |
| pHD-pduC* | Homologous recombination donor vector | Contains homologous regions to insert point mutations in *pduC* (PduC S301A Q336A) through genome editing. Used in BW Pdu to generate BW Pdu*. | This study |
| pTarget-pduC | gRNA expressing vector | gRNA targeting the *pduC* gene of the *pdu* operon for CRISPR/Cas9–mediated genome editing | This study |
| pHD-ΔpduPQ | Homologous recombination donor vector | Contains homologous regions to delete the *pduPQ* genes through genome editing. Used in BW Pdu to generate BW Pdu ΔpduPQ and in BW $Gly^-$ to generate BW $Gly^-$ Pdu. | This study |
| pTarget-ΔpduPQ | gRNA expressing vector | gRNA targeting the *pduP* gene of the *pdu* operon for CRISPR/Cas9–mediated genome editing | This study |

**Table S1: Plasmids used for genome editing in the study.** Their complete sequences are available in the ZENODO repository.

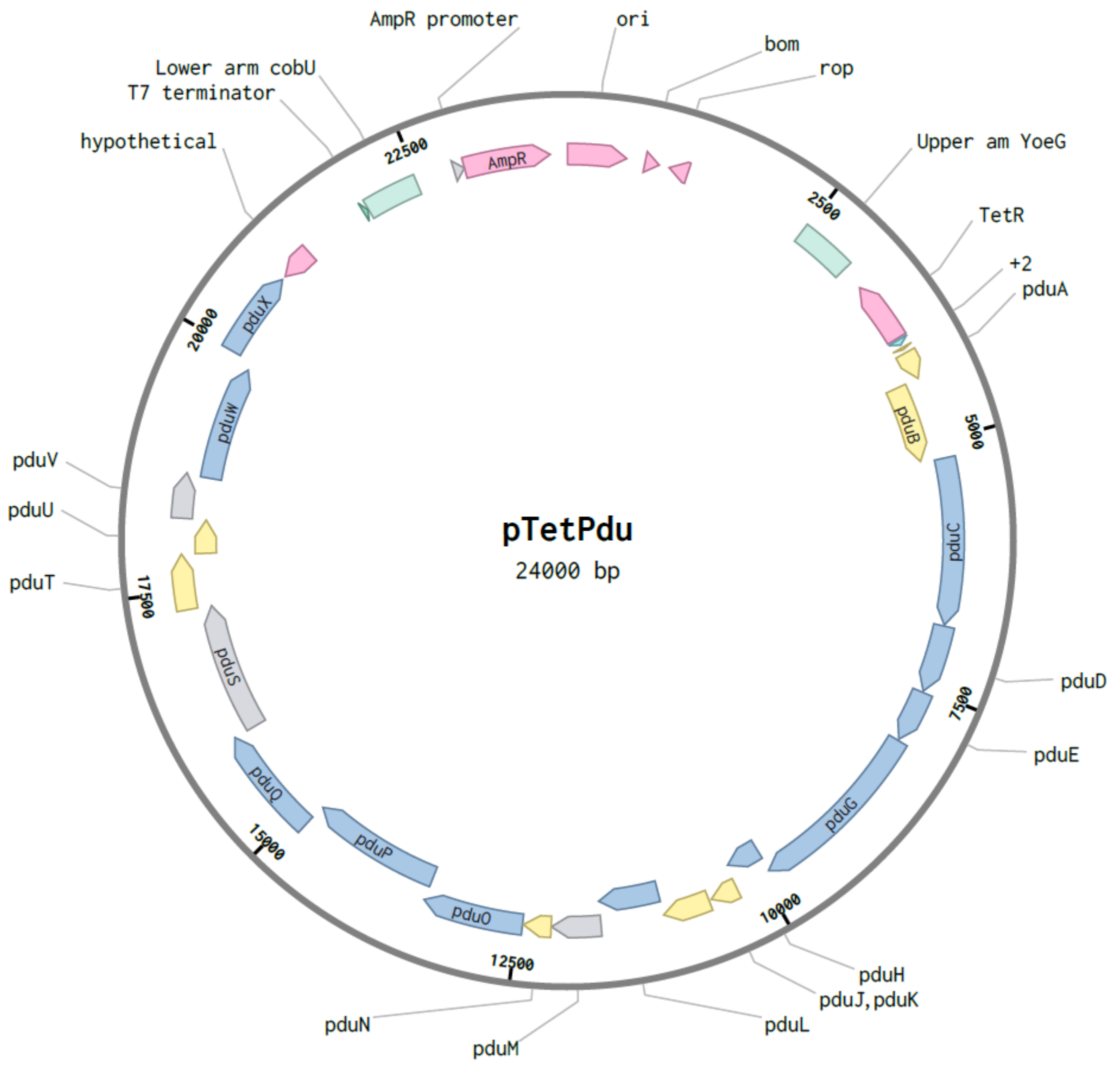


**Figure S1: Map of the pTetPdu plasmid.** pTetPdu (Amp$^R$, ColE1 origin of replication) contains the *pdu* operon of *Citrobacter freundii* cloned under control of the $P_{tet}$ promoter and followed by the $T_{T7}$ terminator. The *pdu* expression cassette is flanked by two homology arms, upper arm *yoeG* and lower arm *cobU*, designed for insertion at the *yoeG-cobU* chromosomal locus of *E. coli* K-12 BW25113 using λ red recombination and a CRISPR-Cas9 based counter-selection strategy[1]. The plasmid map was generated using Benchling (Retrieved from https://benchling.com). The plasmid sequence is available in the ZENODO repository.

[1]Jiang, Y. *et al.* Multigene Editing in the Escherichia coli Genome via the CRISPR-Cas9 System. *Appl Environ Microbiol* **81**, 2506–2514 (2015).

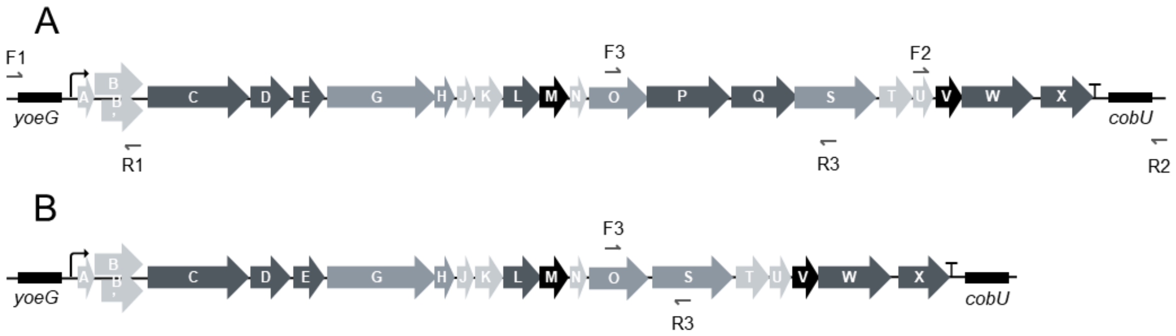


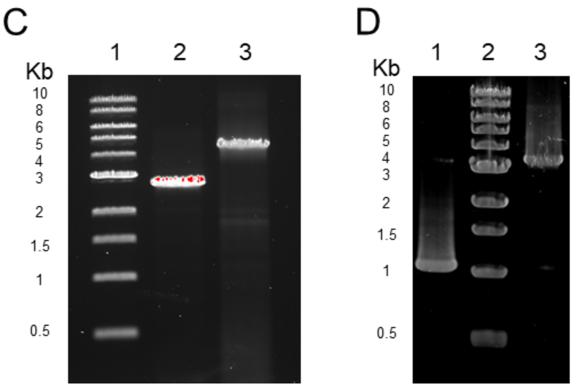


E

| Name | Sequence |
|---|---|
| F1 | AGGTCCGGTTAATTGCTGATACT |
| R1 | TCAGATGTAGGACGGACGATC |
| F2 | CAATCATCGCCTGTGATATCGCCAC |
| R2 | AATCCCCCGGTCATCAGCACCA |
| F3 | ATCAATCGCCTCTCCGACTG |
| R3 | ATTAACGGTCAGGGTACGGC |

**Figure S2: PCRs to verify *pdu* integration and *pduPQ* deletion**. (A) Primer's localisation to amplify the 5' and 3' ends of the integrated *pdu* operon in BW Pdu. (B) Primer's localisation to amplify the locus between *pduO* and *pduS* in BW Pdu *ΔpduPQ*. (C) Agarose gel electrophoresis of the generated amplicons. Lane 1: 1 kb ladder (NEB); Lane 2: 5' PCR product amplified using primers F1/R1 on BW Pdu (expected size: 2.9 kb); Lane 3: 3' PCR product amplified using primers F2/R2 on BW Pdu (expected size 4.7kb). (D) Agarose gel electrophoresis of the generated amplicons. Lane 2: 1 kb ladder (NEB); Lane 1: PCR product amplified using primers F3/R3 on BW Pdu Δ*pduPQ* (expected size: 1.2 kb); Lane 3: PCR product amplified using primers F3/R3 on BW Pdu (expected size 3.5 kb). (E) Primers sequences.

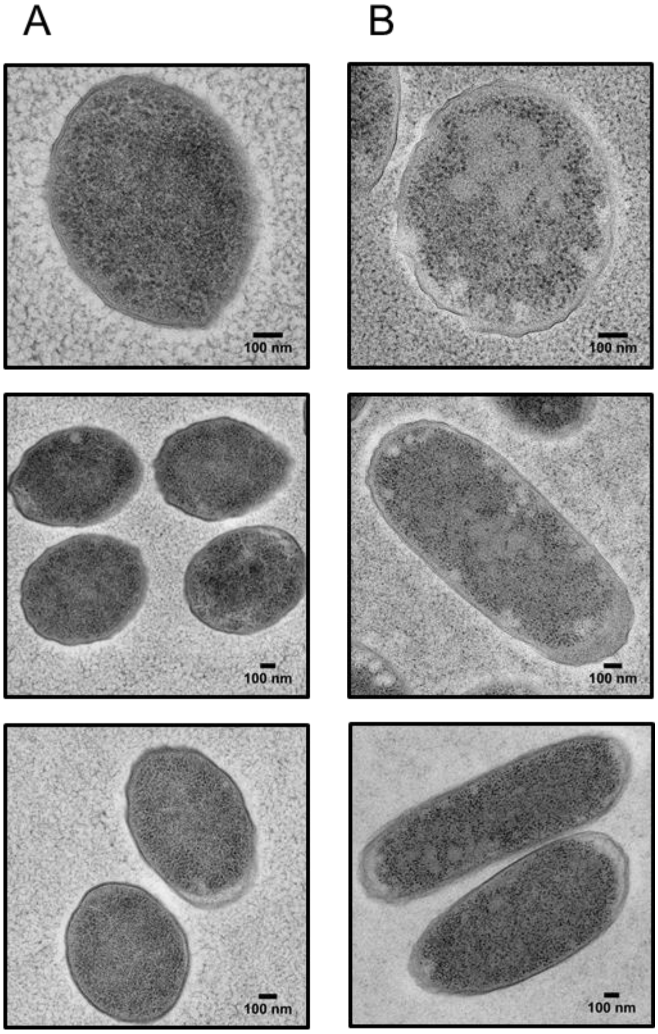


**Figure S3: Additional transmission electron microscopy images** of the (A) BW WT and (B) BW Pdu strains grown aerobically in M9 minimal medium containing D-glucose, 1,2-PDO, cyanocobalamin and tetracycline.

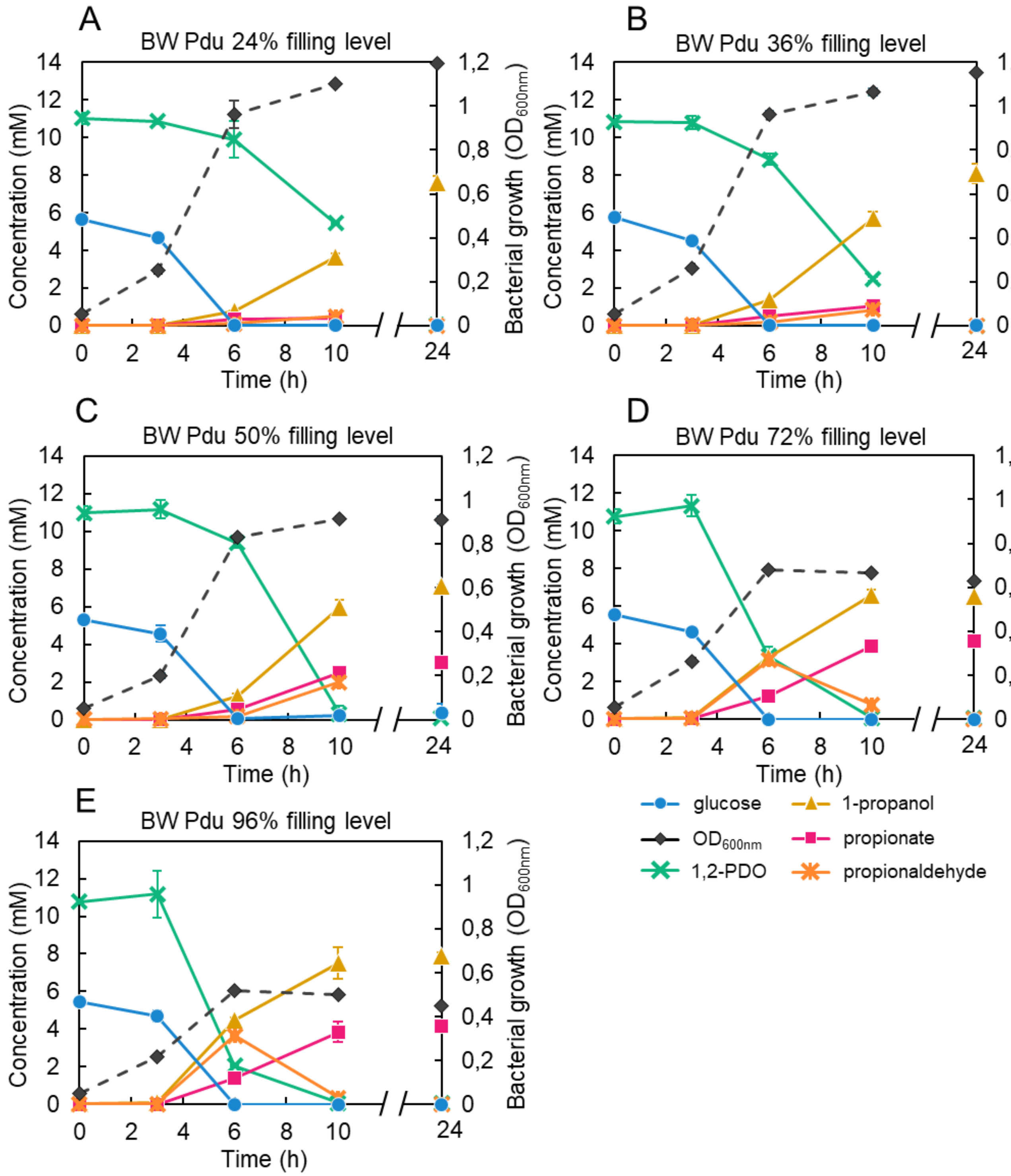


**Figure S4: Effects of the crimp-sealed vials filling level on 1,2-PDO conversion in BW Pdu.** (A) 24%. (B) 36%. (C) 50%. (D) 72%. (E) 96%. Substrate consumption (glucose and 1,2-PDO) and product formation (1-propanol, propionate and propionaldehyde) were quantified using $^{1}$H-NMR. Data are the averages of n = 3 biological replicates, error bars show standard deviations.

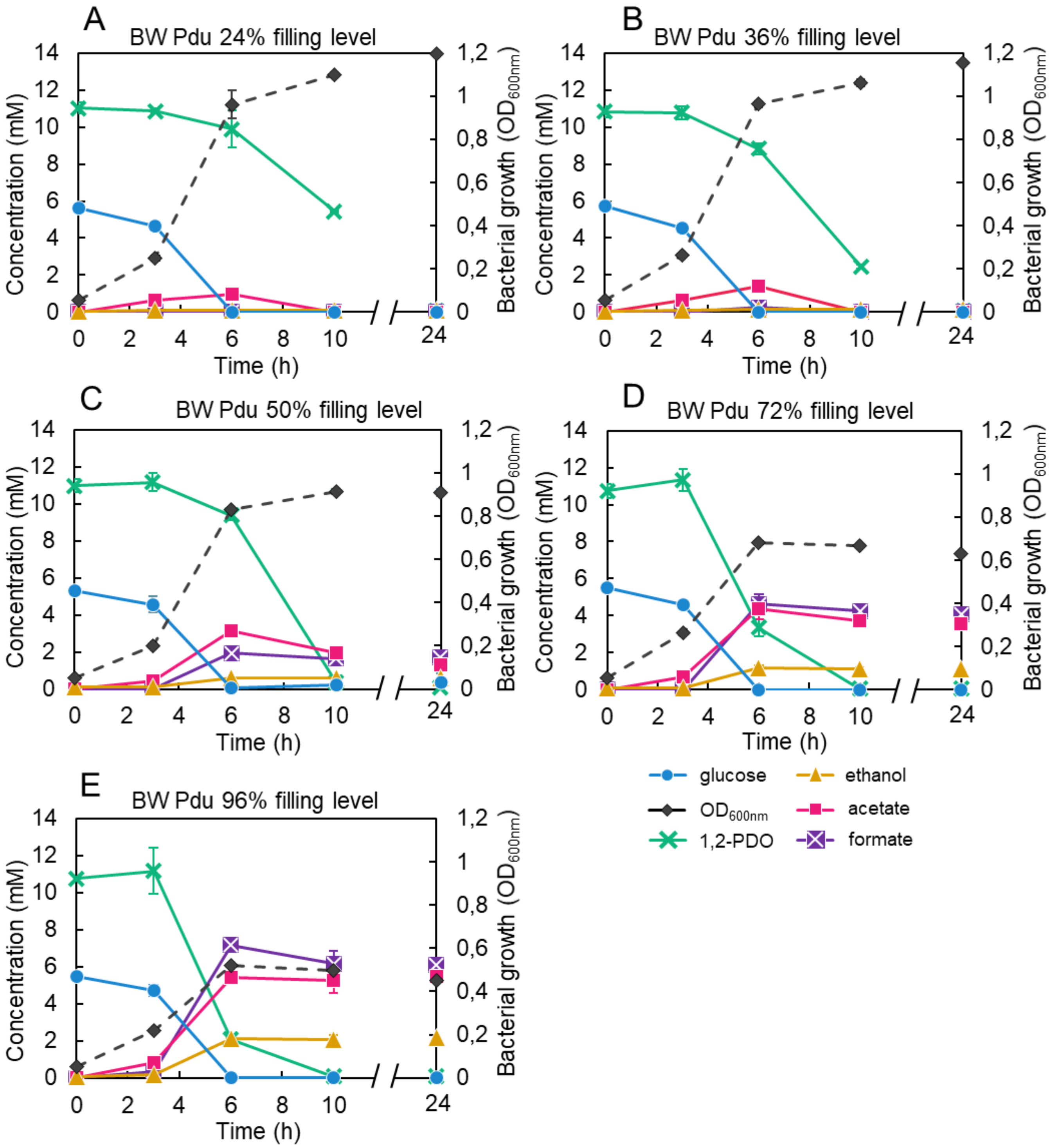


**Figure S5: Effects of the crimp-sealed vials filling level on fermentation products formation in BW Pdu.** (A) 24%. (B) 36%. (C) 50%. (D) 72%. (E) 96%. Substrate consumption (glucose and 1,2-PDO) and product formation (formate, acetate and ethanol) were quantified using $^{1}$H-NMR. These are the same cultures as shown on Figure S4; the fermentation products are displayed here and not in Fig. S4 for clarity. Data are the averages of n = 3 biological replicates, error bars show standard deviations.

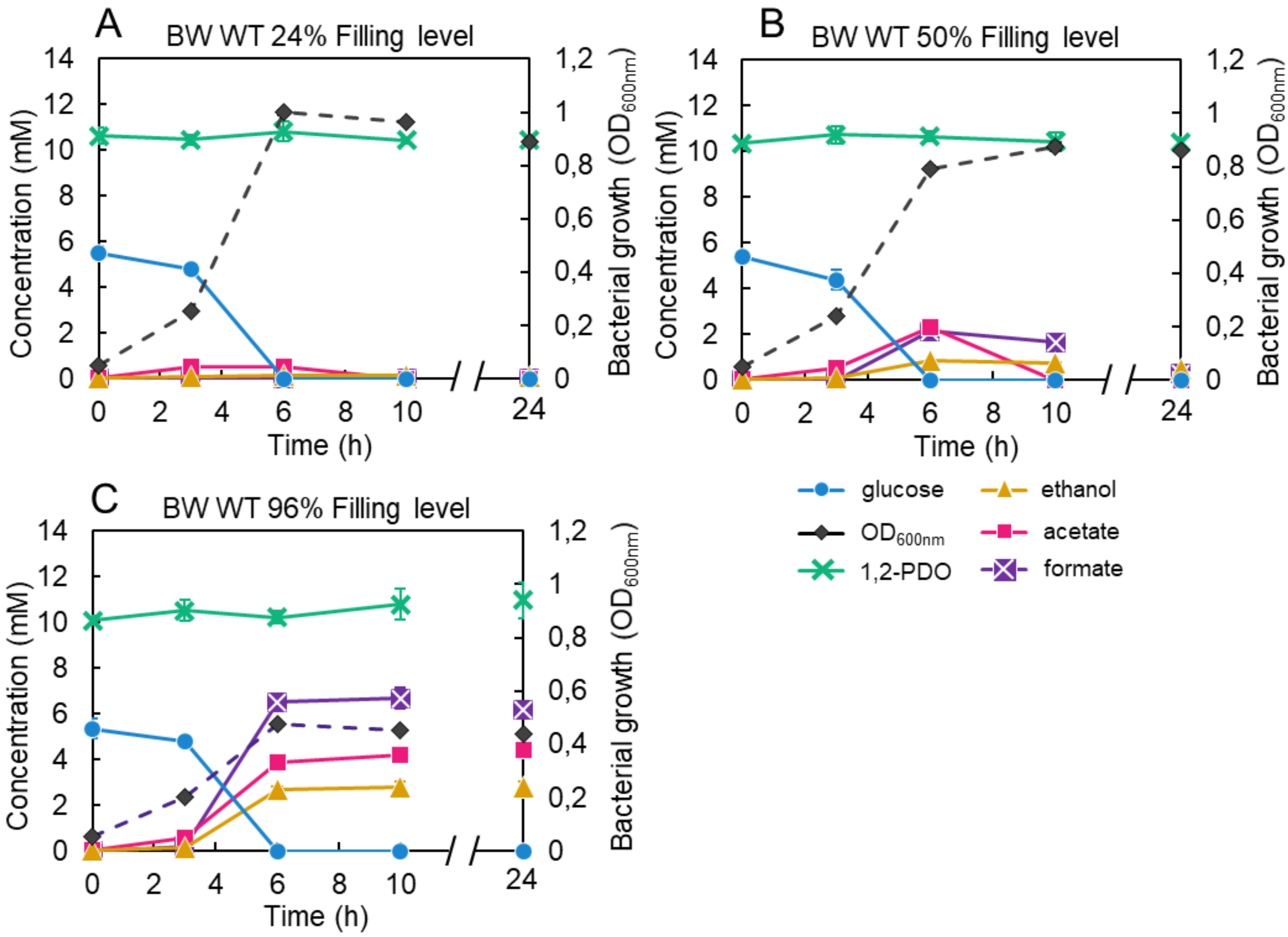


**Figure S6: Effects of the crimp-sealed vials filling level on fermentation products formation in BW WT.** (A) 24%. (B) 50%. (C) 96%. Substrate consumption (glucose and 1,2-PDO) and fermentation product formation (ethanol, acetate and formate) were quantified using $^{1}$H-NMR. Data are the averages of n = 3 biological replicates, error bars show standard deviations.

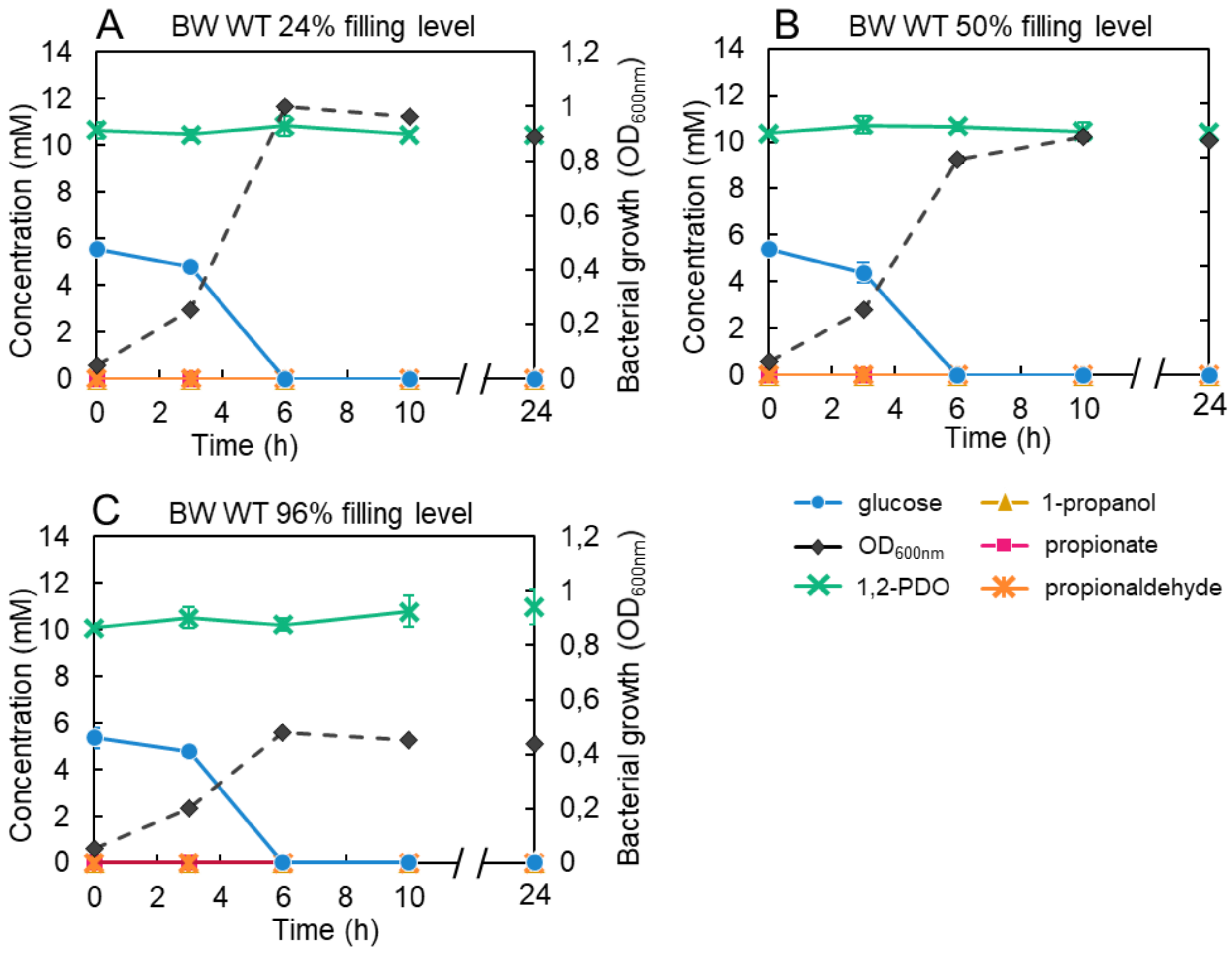


**Figure S7 Effects of the crimp-sealed vials filling level on 1,2-PDO conversion in BW WT.** Various crimp-sealed vial filling levels were used. (A) 24%. (B) 50%. (C) 96%. Substrate consumption (glucose and 1,2-PDO) and product formation (1-propanol, propionate and propionaldehyde) were quantified using $^{1}$H-NMR. These are the same cultures as shown on Figure S6; 1-propanol, propionate and propionaldehyde production are displayed here and not in Fig. S6 for clarity. Data are the averages of n = 3 biological replicates, error bars show standard deviations.

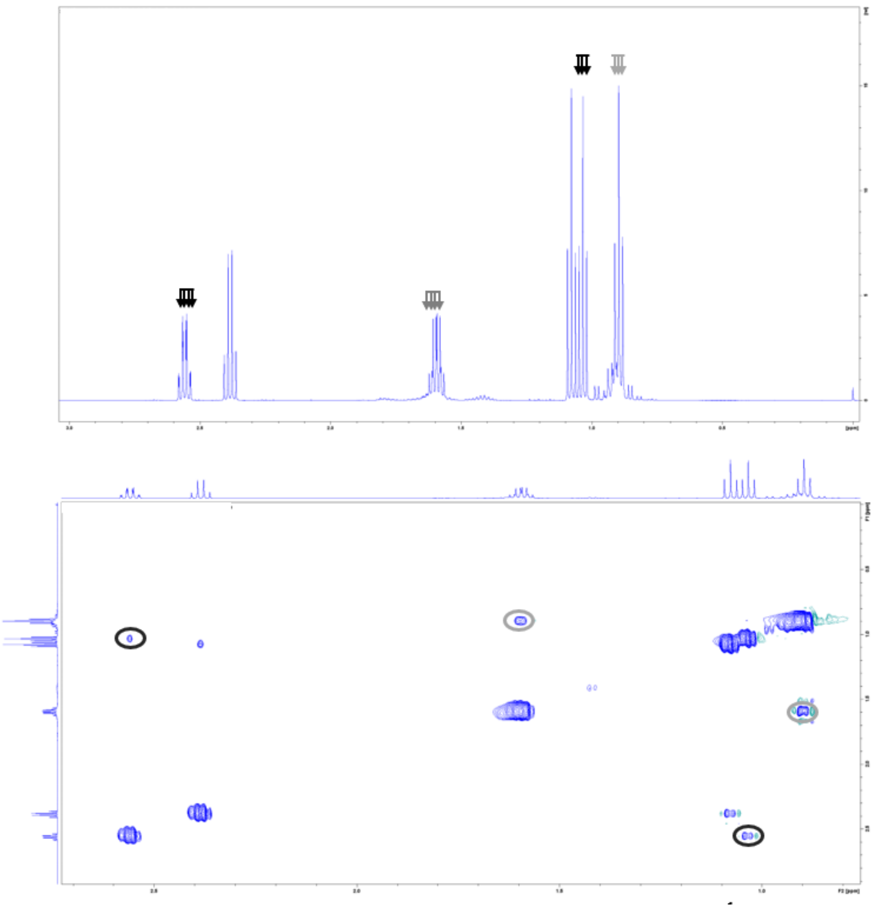

**Figure S8: Identification and quantification of propionaldehyde by $^{1}$H-NMR.** A commercial propionaldehyde solution was diluted to the final concentration of 100 mM in MQ H2O and analysed by NMR at 13°C. (A) One-dimension proton NMR (1H-NMR) experiment; black arrows show peaks associated to propionaldehyde while grey arrows show peaks associated to its hydrated form, 1,1-propanediol. (B) Two-dimensions Correlated Spectroscopy (COSY) experiment to identify protons with spin-spin coupling; the black circle shows correlation in propionaldehyde while the grey one shows correlation in 1,1-propanediol. Propionaldehyde forms an equilibrium of various chemical species in aqueous solution, i.e. spontaneously yielding 1,1-propanediol through hydration or 2,4,6-triethyl-1,3,5-trioxane through polymerization[2]. We identified multiple signals associated to propionaldehyde itself in the 1H-NMR spectrum (A, black arrows): a triplet at 1.04 ppm (3 H+), a multiplet at 2.54 ppm (2 H+) and a triplet at 9.68 ppm (1 H+). These signals were well correlated based on the COSY experiment (B, black circles). Similarly, we identified two correlated signals associated to 1,1-propanediol in the 1H-NMR spectrum (A, grey arrows; B, grey circles): a triplet at 0.90 ppm (3H+) and a multiplet at 1.59 ppm (2H+). Note that some contaminating propionate was also present in the propionaldehyde standard used here. Only trace amounts of trioxane were detected. Consequently, the total concentration of propionaldehyde in solution is calculated as the sum of the propionaldehyde and 1,1-propanediol concentrations.

[2]Corrochano P, García-Río L, Poblete FJ, Rodríguez-Dafonte P. 2010. Spontaneous cyclo-trimerization of propionaldehyde in aqueous solution. Tetrahedron Letters 51:1761–1765.

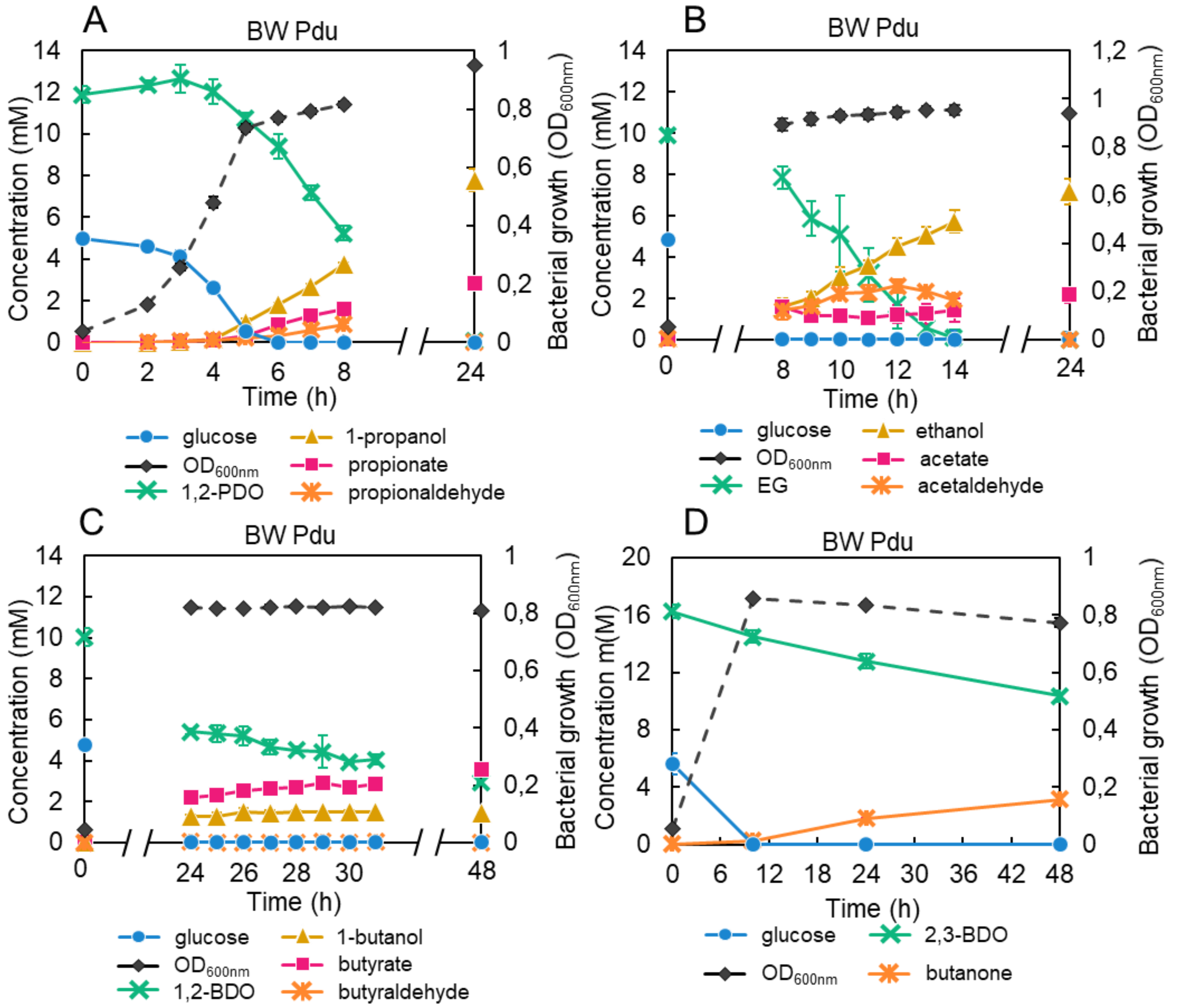


**Figure S9: Substrate consumption and product formation by the BW Pdu strain grown in the presence of different diols at 50% filling level.** (A) 1,2-propanediol (1,2-PDO). (B) Ethylene glycol (EG). (C) 1,2-butanediol (1,2-BDO). (D) 2,3-butanediol (2,3-BDO). Bacterial growth was monitored by measuring $OD_{600}$. Substrate consumption and product formation were quantified using $^1$H-NMR. Data are the averages of n = 3 biological replicates for (A) to (C) and n = 2 biological replicates for (D), error bars show standard deviations.

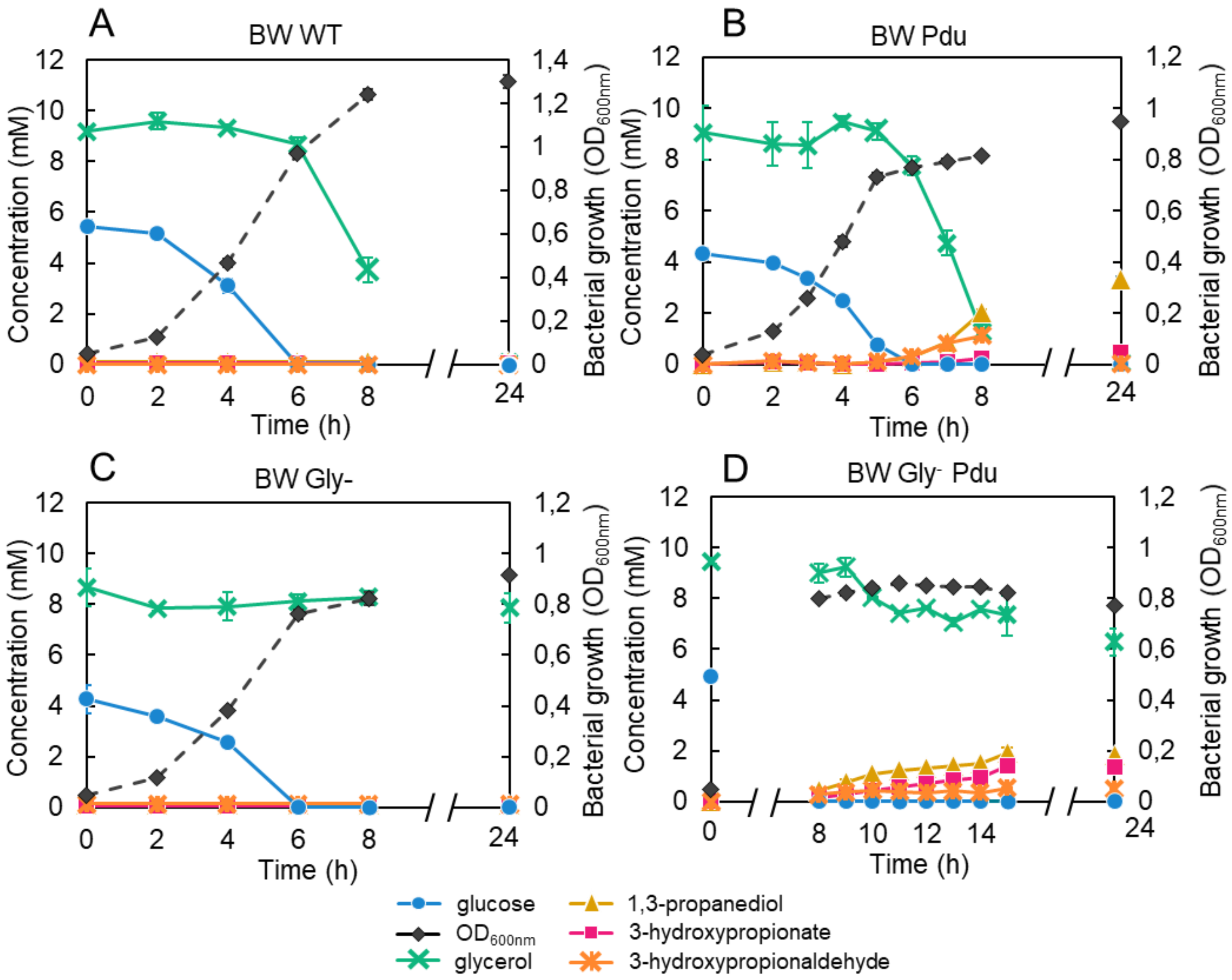


**Figure S10: Glycerol consumption and 1,3-propanediol (1,3-PDO) / 3-hydroxypropionate (3-HP) production at 50% filling level** by the (A) BW WT, (B) BW Pdu, (C) BW Gly$^-$ and (D) BW Gly$^-$ Pdu strains. Bacterial growth was monitored by measuring $OD_{600}$. Substrate consumption (glucose and glycerol) and product formation (1,3-PDO and 3-HP) were quantified using $^1$H-NMR. Data are the averages of n = 3 biological replicates, error bars show standard deviations.

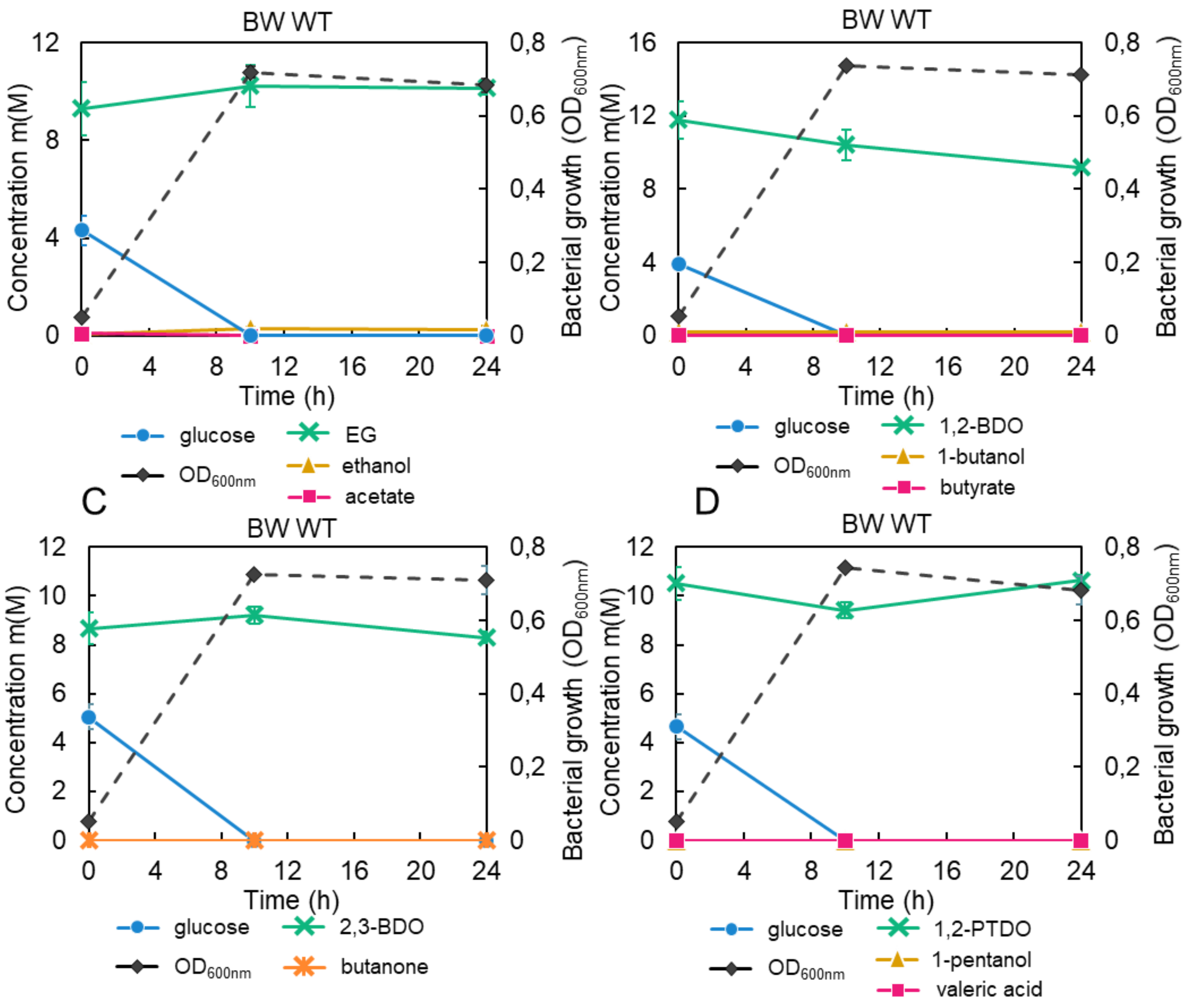


**Figure S11: Substrate consumption and product formation by the BW WT strain grown in the presence of different diols at 50% filling level.** (A) Ethylene glycol (EG). (B) 1,2-butanediol (1,2-BDO). (C) 2,3-butanediol (2,3-BDO). (D) 1,2-pentanediol (1,2-PTDO). Bacterial growth was monitored by measuring $OD_{600}$. Substrate consumption and product formation were quantified using $^1$H-NMR. Data are the averages of n = 2 biological replicates, error bars show standard deviations.

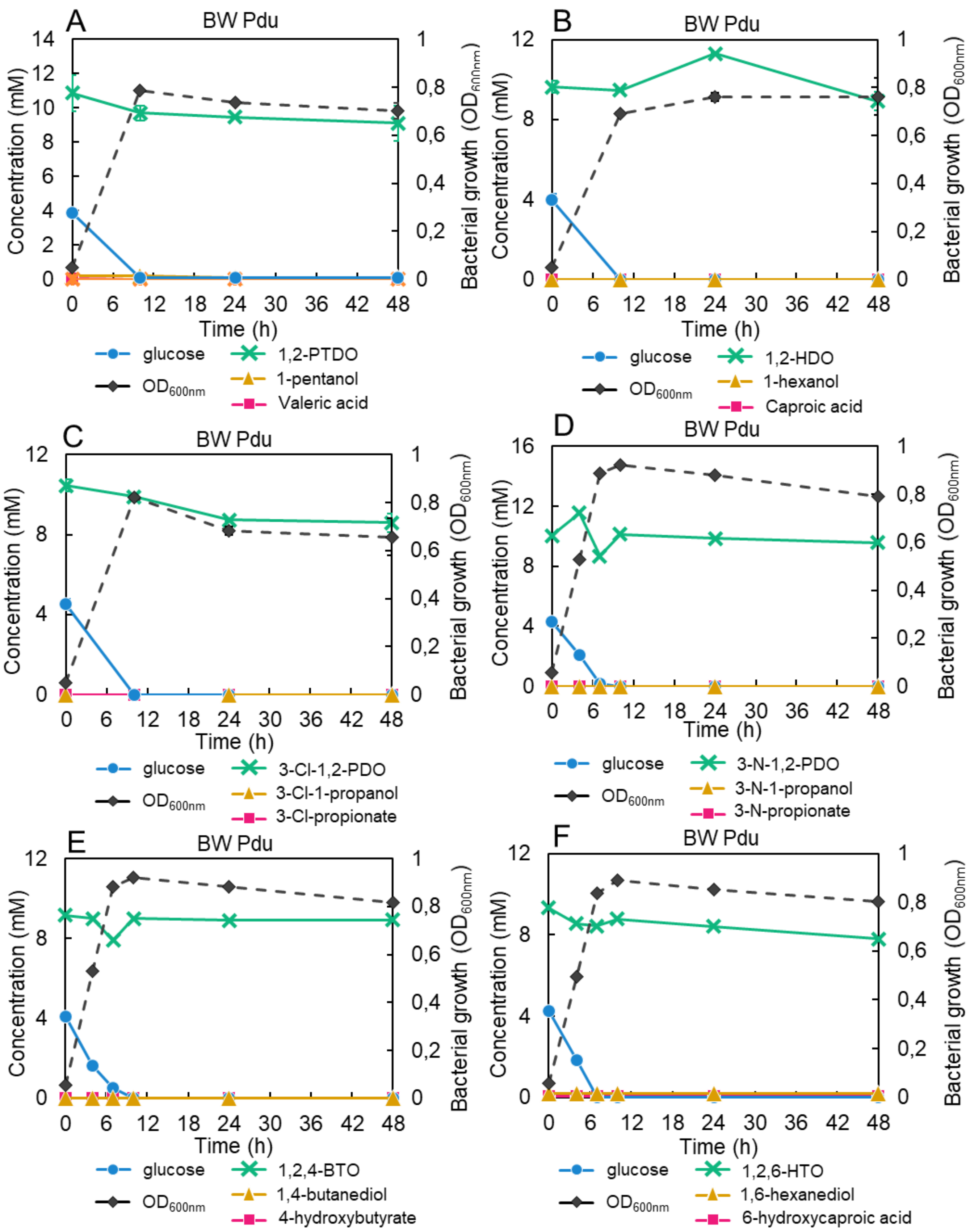
A
BW Pdu
Concentration (mM)
Bacterial growth (OD600nm)
Time (h)
glucose
OD600nm
1,2-PTDO
1-pentanol
Valeric acid
B
BW Pdu
glucose
OD600nm
1,2-HDO
1-hexanol
Caproic acid
C
BW Pdu
glucose
OD600nm
3-Cl-1,2-PDO
3-Cl-1-propanol
3-Cl-propionate
D
BW Pdu
glucose
OD600nm
3-N-1,2-PDO
3-N-1-propanol
3-N-propionate
E
BW Pdu
glucose
OD600nm
1,2,4-BTO
1,4-butanediol
4-hydroxybutyrate
F
BW Pdu
glucose
OD600nm
1,2,6-HTO
1,6-hexanediol
6-hydroxycaproic acid

**Figure S12: Substrates not consumed by the BW Pdu strain.** (A) 1,2-pentanediol (1,2-PTDO). (B) 1,2-hexanediol (1,2-HDO). (C) 3-Cl-1,2-propanediol (3-Cl-1,2-PDO). (D) 3-N-1,2-propanediol (3-N-1,2-PDO). (E) 1,2,4-butanetriol (1,2,4-BTO). (F) 1,2,6-hexanetriol (1,2,6-HTO). Bacterial growth was monitored by measuring $OD_{600}$. Substrate consumption and product formation were quantified using $^1$H-NMR. The crimped-seal vials filling level was 50%. Data are the averages of n = 2 biological replicates for (A) and n = 1 biological replicate for (B) and (F), error bars show standard deviations.

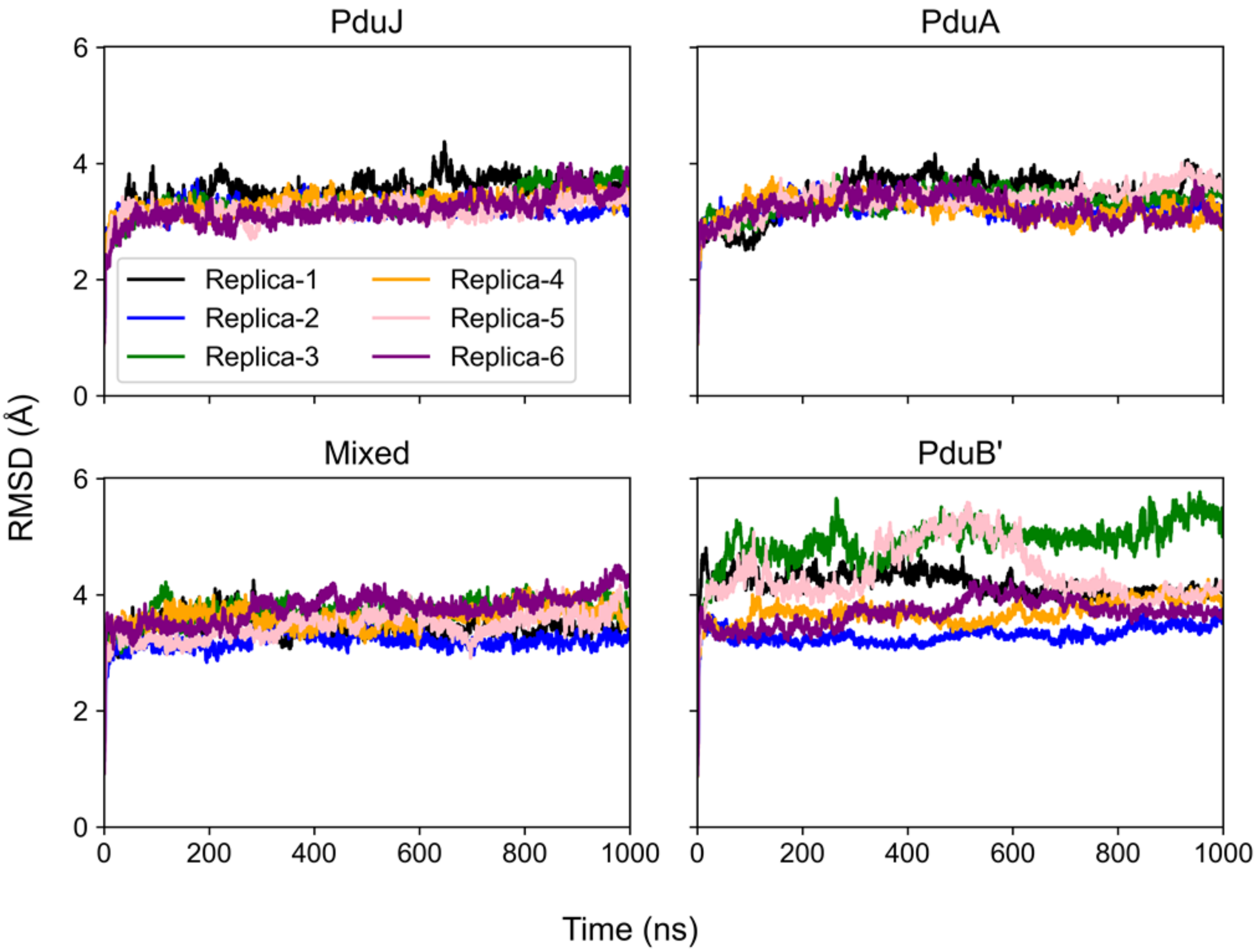


**Figure S13: Root mean squared deviation (RMSD) timeseries for the six simulation replicates with 1,2-PDO present.** The RMSD is calculated on a protein-tile basis, with typically small deviations from the initial structure. Together with visual inspection, this highlights that the proteins remain intact throughout the simulation. To reduce noise, these lines have been smoothed to report a moving average over 20 datapoints separated by 100 ps.

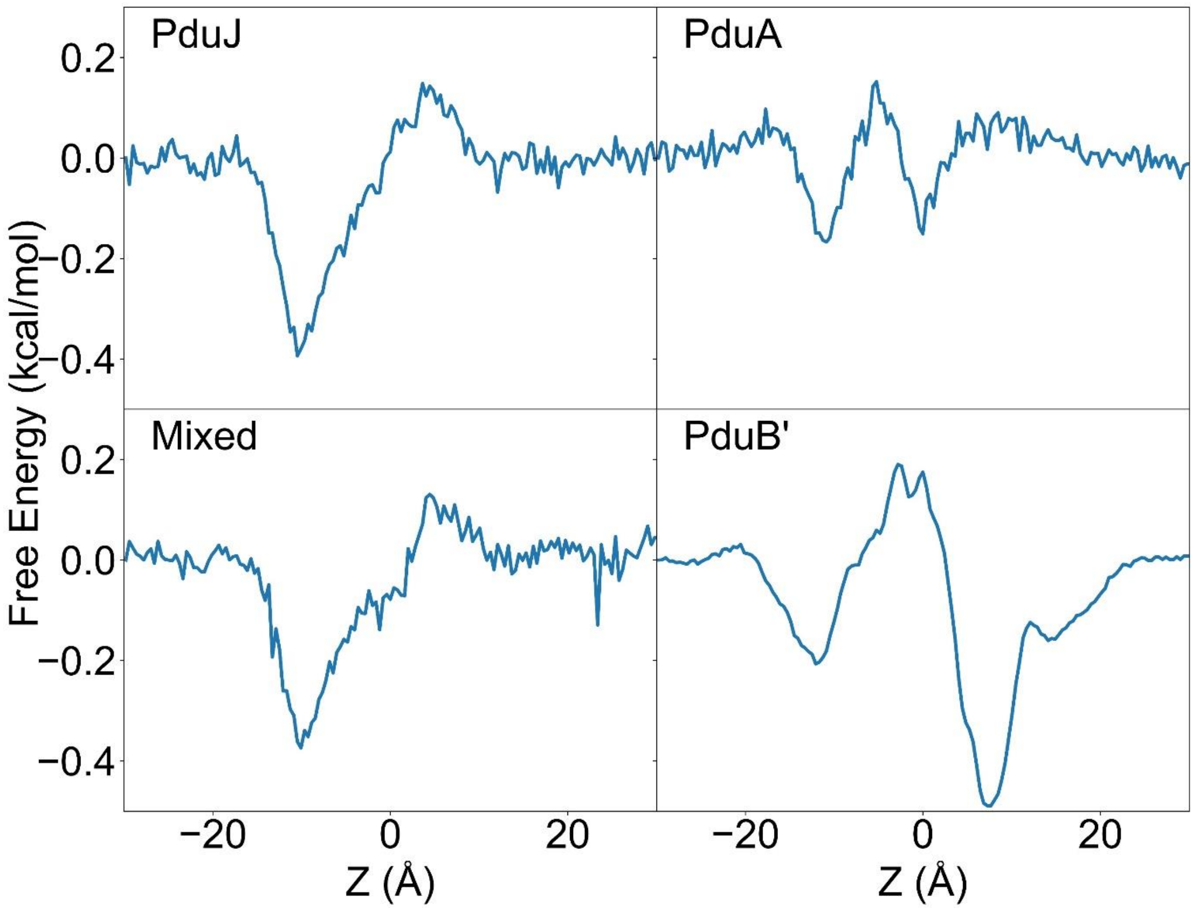


**Figure S14: 1-D Potential of Mean Force (PMF) for each protein-tile for the diffusion of 1,2-PDO.** PMF values were calculated by assessing the free energy across all Z-values at a tiles pore location. Pores of the hexameric tiles, PduJ, PduA, and Mixed, span between r = 0 Å and r = 2.5 Å . For PduB’, the trimeric tile, the three pores span between r = 11 Å and r = 13 Å.

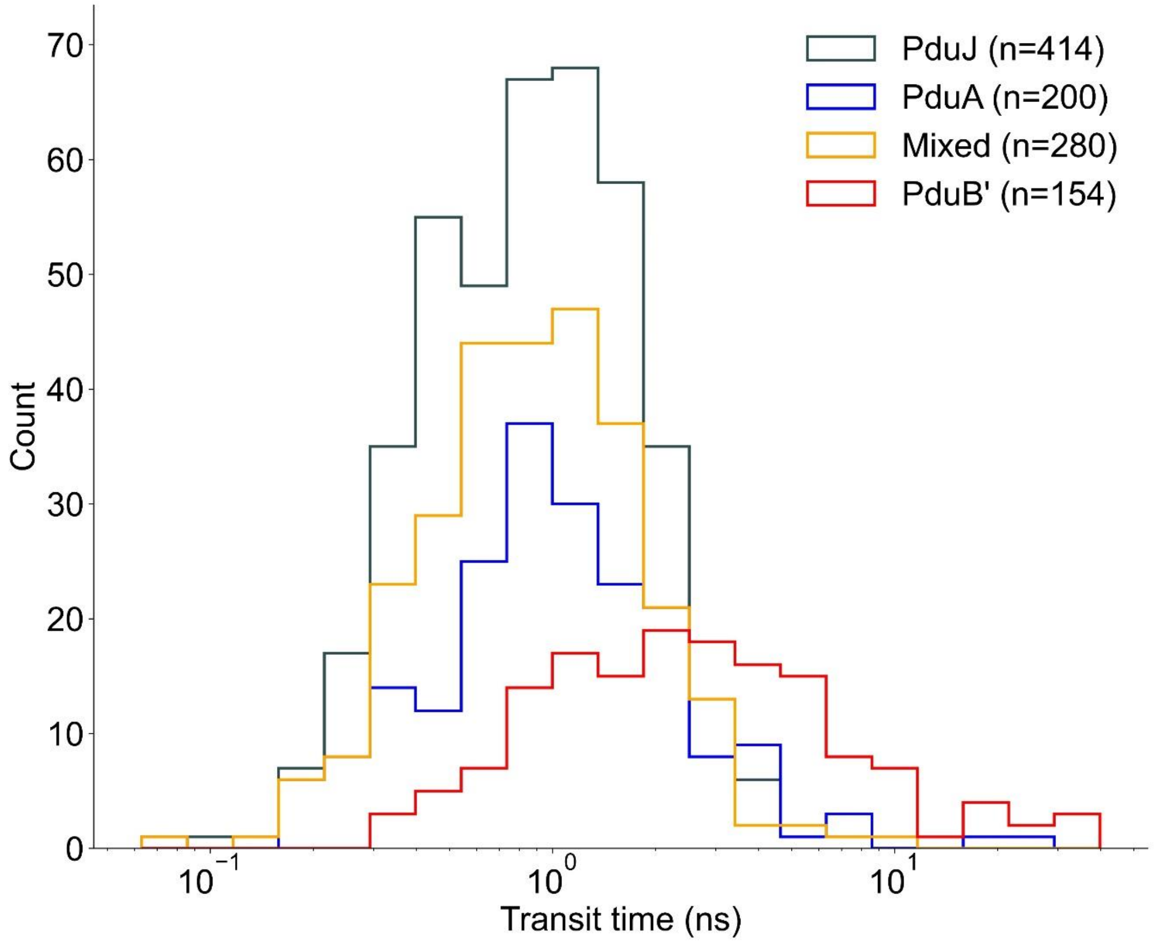


**Figure S15: Distribution of mean transit times for 1,2-PDO molecules across the protein shell pores.** Transit times are defined as the time in which a 1,2-PDO moves from a position of z = 18 Å to z = -18 Å, or in the reverse direction, through a defined pore region. Transit events are accumulated from 6 µs of sampling. Times are represented on a log scale.

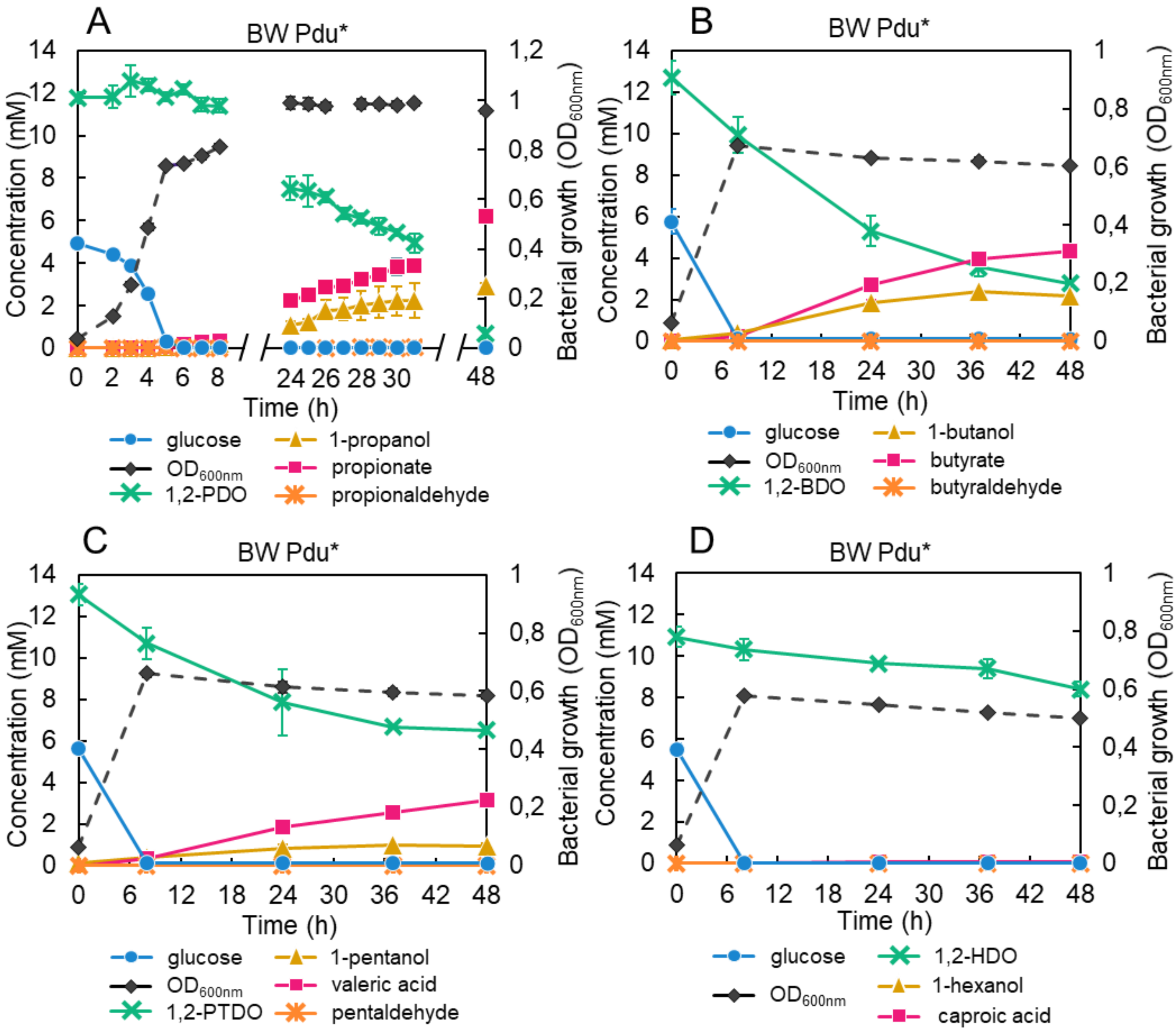


**Figure S16: Substrate consumption and product formation by the BW Pdu* strain grown in the presence of different diols at 50% filling level.** (A) 1,2-propanediol (1,2-PDO). (B) 1,2-butanediol (1,2-BDO). (C) 1,2-pentanediol (1,2-PTDO). (D) 1,2-hexanediol (1,2-HDO). Bacterial growth was monitored by measuring $OD_{600}$. Substrate consumption and product formation were quantified using $^1$H-NMR. Data are the averages of n = 3 biological replicates for (A) and n = 2 biological replicates for (B) to (D), error bars show standard deviations.

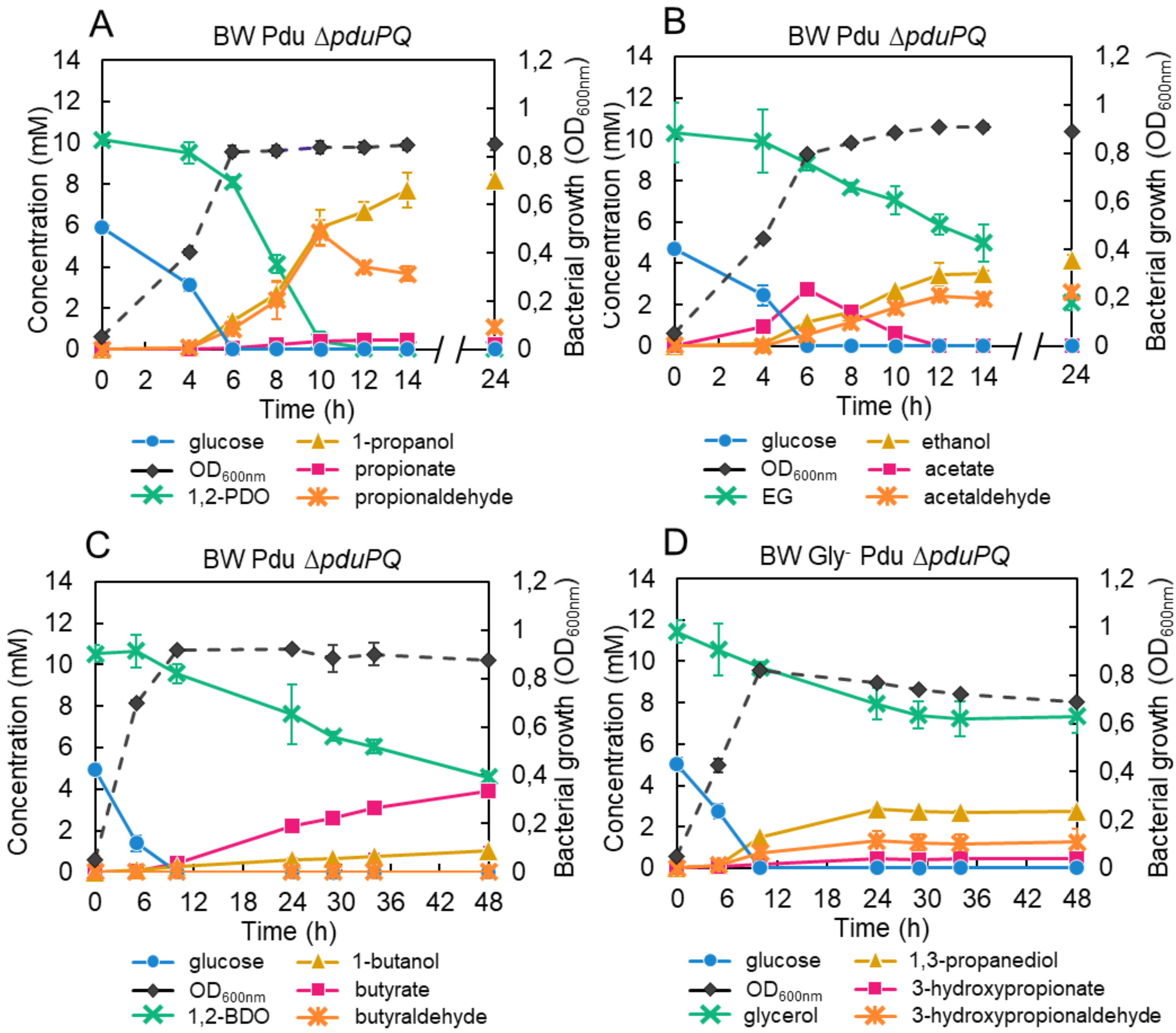


**Figure S17: Substrate consumption and product formation by Δ*pduPQ* mutants in presence of different polyols at 50% filling level.** The BW Pdu Δ*pduPQ* strain was grown in presence of (A) 1,2-propanediol (1,2-PDO). (B) Ethylene glycol (EG). (C) 1,2-butanediol (1,2-BDO). The BW Gly$^-$ Pdu Δ*pduPQ* strain was grown in presence of (D) glycerol. Bacterial growth was monitored by measuring $OD_{600}$. Substrate consumption and product formation were quantified using $^1$H-NMR. Data are the averages of n = 3 biological replicates, error bars show standard deviations.

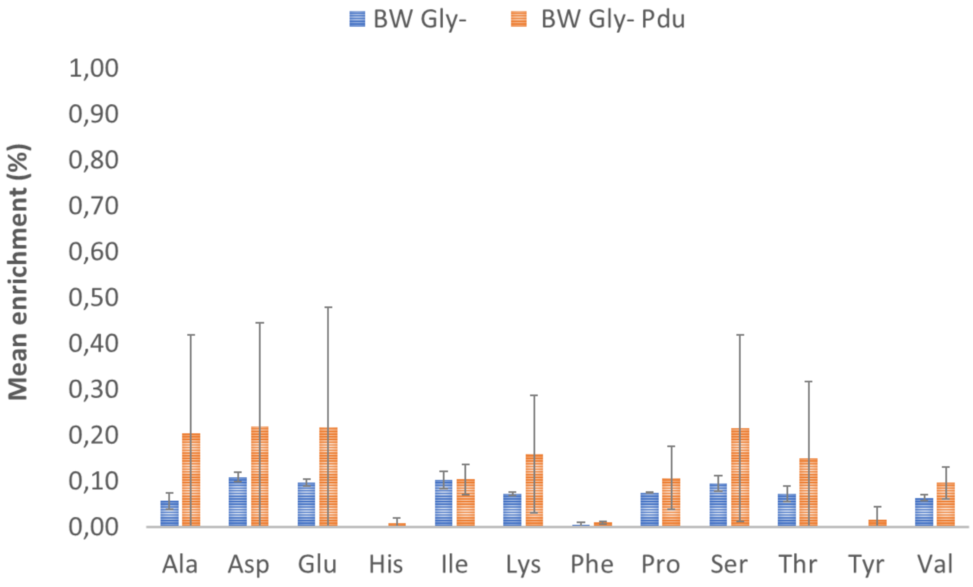


**Figure S18: Mean enrichment detected in the proteinogenic amino acids of the BW Gly$^-$ and BW Gly$^-$ Pdu strains.** Cells were cultivated in a glucose + $^{13}$C-glycerol medium. $^{13}$C-labeling in proteinogenic amino acids was analyzed after 48h of cultivation by mass spectrometry. Data are the averages of n = 3 biological replicates and error bars show standard deviations.
