## Supplemental Material and Methods for "Small but Mighty: The Surprising Metabolic Range of a Bacterial Microcompartment"

### *PCR amplification*

Colony PCR was performed using the OneTaq® DNA Polymerase kit (New England Biolabs). A small amount of a bacterial colony grown on selective agar plates was picked with a sterile toothpick and directly transferred into a 25 µL PCR reaction mixture containing 1× OneTaq Standard Reaction Buffer, 200 µM of each dNTP, 0.2 µM of each primer, and 0.5 U of OneTaq DNA Polymerase. Nuclease-free water was added to reach the final reaction volume. PCR amplification was performed using the following thermal cycling conditions: initial denaturation at 94°C for 30 s, followed by 30 cycles of denaturation at 94°C for 15 s, annealing at 60°C for 30 s, and extension at 68°C for 1 min per kb of expected product. A final extension step was performed at 68°C for 5 min. Amplification products were analyzed by electrophoresis on a 1% agarose gel prepared in 1× TAE buffer and stained with ethidium bromide**.**

### *BMC purification*

Pdu BMCs were purified by differential centrifugation following a previously described method [1]. Briefly, 400 mL bacterial cultures supplemented with vitamin B12 (200 nM) and anhydrotetracycline (80 ng/mL) as an inducer of the heterologous *pdu* operon were harvested by centrifugation at 4,600 × g for 5 min at room temperature. The resulting cell pellets were resuspended in 20 mL of buffer A (100 mM potassium phosphate buffer [pH 8.0], 500 mM KCl, and 12.5 mM $MgCl_2$). Cells were lysed by resuspending the pellets in 20 mL of lysis buffer (lysozyme 1 mg/mL, β-mercaptoethanol 5 mM, DNase I 25 U/mL, AEBSF 0.4 mM, and BugBuster reagent 50% v/v) in water. The suspension was incubated for 30 min at room temperature with 60 rpm of agitation to allow cell lysis. Cell debris were removed by centrifugation at 12,000 g for 5 min at 4°C and the supernatant was subjected to a second clarification step under the same conditions. The clarified lysate was then centrifuged at 21,000 × g for 20 min at 4°C to pellet the Pdu BMCs. The resulting pellet was washed by resuspension in buffer A supplemented with BugBuster reagent and 0.4 mM AEBSF, followed by centrifugation at 21,000 × g for 20 min at 4°C. The pellet was resuspended in 1 mL of buffer B (100 mM potassium phosphate buffer [pH 8.0], 50 mM KCl, and 5 mM $MgCl_2$). The suspension was subsequently clarified by three successive centrifugation steps at 12,000 g for 1 min, transferring the supernatant to a fresh tube after each step. The final supernatant containing purified Pdu BMCs was stored at 4°C until further use. A bicinchoninic acid assay was used to measure the concentration of the purified Pdu BMCs. Samples for SDS-PAGE analysis were prepared by mixing 6 µg of the generated protein fraction with 4X Laemmli buffer containing 10% (v/v) β-mercaptoethanol. Samples were denaturated at 95°C for 10 min and then deposited into an Any kD Mini-PROTEAN TGX Precast Protein Gel (Biorad), with Precision Plus Protein ladder (Biorad) in a separate well. Migration was performed at 60 V for 40 min in a Tris-Glycine-SDS (TGS) running buffer. Protein bands were revealed employing InstantBlue (Abcam).

#### *High pressure freezing and transmission electron microscopy*

BW WT and BW Pdu cultures grown aerobically in M9 medium containing D-glucose, 1,2-PDO, cyanocobalamin as well as tetracycline were harvested during exponential phase (samples of 1.25 mL at $OD_{600nm}$ = 0.8). After a centrifugation step at 4000 × g, 3 min, room temperature, supernatants were removed and pellets resuspended in 1 ml 100 mM PBS buffer pH 7. This washing was repeated twice more and supernatants were eventually discarded while keeping pellets slightly wet. For cryofixation, bacteria were placed between 3 mm gold-coated copper carriers (types A and B, 200 µm deep, Leica Microsystems). The sandwiched samples were frozen using a Leica EM ICE high -pressure freezer (Leica Microsystems, Austria). After freezing, the samples were immersed in a substitution medium consisting of 2% osmium tetroxide, 0.5% glutaraldehyde, and 0.1% uranyl acetate in absolute acetone, and pre-cooled to -90 ◦C using an Automated Freeze-substitution System (Leica Microsystems). The AFS program was run as follows: −90°C for 57 h, −90°C to −60°C for 6 h (5°C/h), −60°C for 8 h, −60°C to −30°C for 6 h (5°C/h), −30°C for 8 h, -30 to 0°C for 6 h (5°C/h), 0 to +15°C for 15min. After freeze-substitution, the samples were washed three times with absolute acetone at room temperature and embedded in EMBed 812 epoxy resin. After 48 h of polymerization at 60 °C, ultrathin sections (80 nm thick) were mounted onto 200-mesh Formvar carbon-coated copper grids. Finally, sections were stained with Uranyless and lead citrate. Grids were examined with a TEM (Jeol JEM-1400, JEOL Inc, Peabody, MA, USA) at 80 kV. Images were acquired using a digital camera (Gatan Rio9, Gatan Inc, Pleasanton, CA, USA).

#### *Deletion of the glycerol utilization pathways*

The λ red recombination method [2] was used to generate the Δ*dhaKLM* Δ*glpK* Δ*gldA* triple knock out strain. The introduced antibiotic resistance cassettes were removed with the FRT/FLP recombination system. All constructs were subsequently verified by colony PCR and Sanger sequencing (Genewiz).

#### *Propionaldehyde analysis.*

The total propionaldehyde concentration in solution, when required, was calculated as the sum of the propionaldehyde and 1,1-propanediol concentrations [3] (Fig.S6). A homonuclear 1H-1H Correlation Spectroscopy (COSY) spectrum was first collected on a 100 mM propionaldehyde standard in aqueous solution. We used the cosyphpr sequence with the following acquisition parameters: 286 K, with 16 dummy scans followed by 4 scans of 512 real data points with a spectral width of 10 ppm in F1 and 4096 real data points with a spectral width of 10 ppm in F2.

The generated 2D spectrum allowed to identify coupled $^{1}H$ nuclei over three bonds in the 1D $^{1}H$-NMR spectrum and to assign peaks to propionaldehyde or 1,1-propanediol, respectively.

### *Isotopic labeling experiment*

Cultures were grown in M9 minimal medium containing glycerol-2-$^{13}C$ (Sigma Aldrich 489484-1G). One milliliter of culture was sampled every 2 h. Optical density (OD) was measured, and samples were centrifuged for 30 s at 16,500 × g to separate the pellet from the supernatant. Cell pellets were then lysed with 250 µL of 6 N HCl at 110 °C for 15 h to hydrolyse peptide bonds between proteinogenic amino acids. HCl was subsequently evaporated under reduced atmospheric pressure (20 mbar at 40 °C). Hydrolysates were then washed twice with distilled water using the same evaporation procedure. Dried hydrolysates were resuspended in 50 µL of distilled water and filtered through 0.45 µm filters. Hydrolysates were finally diluted proportionally with distilled water according to OD for analysis as follows: for a sample with OD = 1, the hydrolysate was diluted 100-fold, whereas for OD = 0.1, the hydrolysate was diluted 10-fold.

Analyses were performed on an LC-MS platform consisting of a Thermo Scientific Vanquish™ Focused UHPLC Plus system coupled to a Thermo Scientific Q Exactive™ Plus hybrid quadrupole-Orbitrap™ mass spectrometer (Thermo Fisher Scientific™, Waltham, MA) equipped with a heated electrospray ionization probe. Chromatographic separation of amino acids was carried out using a SUPELCO PFP column (150 × 2.1 mm I.D., 5 µm) and an HS F5 guard column (5 µm, 20 × 2.1 mm). The column was maintained at 30 °C and the flow rate was set to 0.25 mL/min. The solvent system consisted of solvent (A), 0.1% formic acid in water, and solvent (B), 0.1% formic acid in acetonitrile. The gradient was adapted from the method described by Boudah et al. Solvent B was modified as follows: 0 min: 2%, 2 min: 2%, 10 min: 5%, 16 min: 35%, 20 min: 100%, and 24 min: 100%. The column was then re-equilibrated for 6 min under initial conditions before injection of the next sample. The injection volume was 5 µL and the autosampler temperature was maintained at 4 °C. Mass detection was performed in positive electrospray ionization (ESI) mode. Mass spectrometer settings were as follows: spray voltage 5.0 kV, capillary and desolvation temperatures of 300 and 320 °C, respectively, and maximum injection time of 100 ms. Nitrogen was used as sheath gas (40 arbitrary units) and auxiliary gas (10 arbitrary units). Automatic gain control (AGC) was set to 1e6, with a resolution of 70,000 over an m/z range of 50–750. MS analyses were performed in Full Scan mode. Data acquisition was carried out using Thermo Scientific Xcalibur software.

$^{13}C$-isotopologue distributions were determined by matching accurate masses (±5 ppm) and retention times using TraceFinder™ 4.1 software (Thermo Fisher Scientific, Waltham, MA, USA). Peak areas corresponding to individual isotopologues were corrected for natural isotope abundance and tracer isotopic purity using IsoCor [4]. The degree of $^{13}C$ enrichment was calculated as the sum of the fractional abundances of each isotopologue multiplied by the number of incorporated $^{13}C$ atoms, divided by the total number of carbon atoms in the metabolite fragment, according to the equation:

$$^{13}C\ enrichment\ (\%) = \frac{\Sigma(\mathrm{M_i} \times \mathrm{i})}{n}$$

Where n is the number of carbon atoms and $M_i$ is the corrected abundance of isotopologue i.

### *Molecular dynamics simulations.*

#### System construction

To determine permeability through the pores found in these BMCs, we develop an atomic simulation model featuring 4 BMC tiles arrayed in a plane. Protein sequences for each component tile were derived from previous sequencing of *Citrobacter freundii*. Hexameric PduA and PduJ tile structures were predicted using Alphafold 3 [5]. A trimer was produced based on the sequences for PduB'. PduB and PduB' share the same sequences, except that PduB possesses a 37 amino acid long N-terminal extension that is not present in PduB'. PduB' was chosen here because the N-terminal extension structure cannot be modeled confidently by Alphafold 3. The final tiling component was a mixed hexamer comprised of three monomers from each of PduA and PduJ with its structure similarly predicted by Alphafold 3. The system was assembled by arranging the protein tiles normal to the z-axis in a rhomboid configuration based on the size of individual tiles, analogous to prior planar BMC simulations [6–8]. Previous work in comparing simulations using bacterial microcompartment shells vs sheets has shown that pore dynamics are highly conserved between the two methods of simulation [7]. Using planar sheets to simulate these pore dynamics comes at a significant computational discount as opposed to shells, as the sheets have fewer atoms to simulate.

The sheet system was then solvated using the VMD 1.9.4a58 solvate package and ionizing using the autoionize package to establish the NaCl concentration at 150 mM [9]. To avoid water filling the interfaces between tiles, water was placed in a narrow and tall box at the center of the system after which the system was equilibrated for 2 nanoseconds with alpha carbons constrained to allow the box to compress and for the water to fill the BMC pores. Finally, a specific metabolite was added to random positions to bring its concentration up to 500 mM and ensure that both sides of the bacterial microcompartment shell are isotonic. The metabolites used were glycerol, ethylene glycol, 1,2-propanediol, 1,2-butanediol, 1,2-pentanediol, 1,2-hexanediol, and 1,2,6-hexanetriol, and each metabolite system was independent with a single metabolite present.

### Simulation Protocols

Each of the 7 molecular systems was simulated six times to collect sufficient molecular transit events to collect robust statistics. Each replicate was 1 µs (42 µs total). The CHARMM36m forcefield for proteins was used as well as the TIP3 water model that aligns with this forcefield to model these systems in NAMD 3.0 [10–12]. The topology for all metabolites was generated by the CHARMM-GUI ligand reader and modeler by submitting SDFs of 3D conformations from PUBCHEM [13]. All metabolites with chiral centers were chosen to be solely in the S conformation for these simulations. For these simulations, a nonbonded switching distance of 10 Å and a cutoff distance of 12 Å were used, as is typical in CHARMM36. Long range electrostatics were treated with particle mesh Ewald with a gridspacing of 1.2 Å [14]. Systems were minimized for 2500 steps with alpha carbons constrained followed by 1,000,000 steps (2 ns) of equilibrium with continued constraints on alpha carbons to avoid significant structural deformation of the proteins during the initial equilibration period. Finally, each replicate was run for 1 µs (6 µs per metabolite, 42 µs total) using a Langevin piston barostat and thermostat to regulate pressure and temperature, respectively [15,16]. The barostat was set as a semi-isotropic ensemble at 1 atm, and the thermostat was set to 298K. All simulations were run using a flexible cell without a constant ratio.

### Analysis

Atomistic molecular dynamics simulations were run of a simplified Pdu BMC tile structure to assess the permeability of EG, 1,2-PDO, 1,2-BDO, 1,2-PTDO, 1,2-HDO, and 1,2,6-HTO.

Simulations were analyzed using python-enabled VMD 1.9.4a58 and are provided in a Zenodo repository along with select simulation inputs and outputs. The NumPy library was utilized for data computation and manipulation along with Matplotlib for visualization [17,18].

To verify the stability of protein structures used in these simulations, root-mean-square-deviation (RMSD) over time was utilized. Figure S10 shows that RMSD only moderately exceeds the resolution of structures solved by x-ray crystallography for BMC hexamers with solved structures [19].

### Determining Permeability through Observed Transition Events

In our simulations, there cannot be an explicit concentration gradient, as the compartments above and below the protein sheet are connected across the periodic boundary. Instead, we compute permeability using time-tested techniques developed for membrane simulations and have since been applied to bacterial microcompartment systems [6,20,21]. In this construction, permeability is measured through the rate of transition events (r) and the concentration of the metabolite in the simulation box ($c_w$) as listed in the following equation:

$$P = \frac{r}{2c_w} = \frac{transition\ events}{2c_w * time}$$

Using this definition of permeability, transition events in both directions were tracked by measuring the position along the membrane normal for each metabolite over the simulation trajectory. Transition events were tabulated and used to calculate permeability values for each metabolite through each pore as listed in (Table 3).

To investigate the structure of individual tile pores, the HOLE program was used [22]. HOLE utilizes a space filling model to find the largest sphere that can fit in standardized points along the z-axis, which, when computed in a frame-by-frame basis, provides a representative pore structure. The HoleHelper Tcl plugin, previously used for an analysis comparing pore structures in shells and sheets, was used to assist in this analysis [7].

**Assessing Metabolite Flux based on Observed Permeability**

The concentration gradient needed to support a given flux can be computed by rearranging Fick's law. To give a concrete example, consider a BMC that can turn over 1 molecule per second at steady state, such that the rate for substrate and product transit is 1 molecule per second. Assuming the BMC to be spherical with a radius of 50nm, the concentration gradient needed to sustain that flux can be computed by the following formulation:

$$J = P \cdot A \cdot \Delta C$$

The imputed concentration gradient is quite small, particularly since these metabolites are being fed to the organism and thus would be available at relatively high concentrations. Thus, we anticipate that enzyme turnover would only slow once the concentration gradient approaches the Michaelis constant, which are often in the micromolar range. The permeability coefficients calculated for the metabolites through PduA are all within an order of magnitude of 0.10 cm*$s^{-1}$. These permeability coefficients would correspond to a flux of tens of thousands of molecules per second, which would imply many active enzymes operating at near peak efficiency.
